# Nociceptor to macrophage communication through CGRP/RAMP1 signaling drives endometriosis-associated pain and lesion growth

**DOI:** 10.1101/2023.08.28.555101

**Authors:** Victor Fattori, Tiago H. Zaninelli, Fernanda S. Rasquel-Oliveira, Olivia K. Heintz, Ashish Jain, Liang Sun, Maya L. Seshan, Daniëlle Peterse, Anne E. Lindholm, Raymond M. Anchan, Waldiceu A. Veri, Michael S. Rogers

## Abstract

Endometriosis is a debilitating and painful gynecological inflammatory disease affecting approximately 15% of women. Current treatments are ineffective for a significant fraction of patients, underscoring the need for new medical therapies with long-term benefits. Given the genetic correlation between migraines and endometriosis, we sought evidence for the role of CGRP-mediated neuroimmune communication in endometriosis. We found that mouse and human endometriosis lesions contained CGRP and RAMP1. In mice, nociceptor ablation reduced pain, monocyte recruitment, and lesion size, suggesting that nociceptors support endometriosis lesions. *In vitro,* CGRP-treated macrophages showed impaired efferocytosis and supported endometrial cell growth in a RAMP1-dependent manner. Treatment with FDA-approved drugs that block CGRP-RAMP1 signaling reduced evoked and spontaneous pain, and lesion size. Since the lack of drug efficacy at reducing ongoing pain drives most endometriosis therapy failure, our data demonstrating effectiveness of non-hormonal and non-opioid CGRP/RAMP1 blocking therapies may lead to clinical benefit for endometriosis patients.

## Introduction

Nociceptors, the afferent sensory neurons that transmit pain or itch, detect environmental cues and potentially damaging stimuli, such as pathogens or injured tissue (Baral et al., 2019; Chiu et al., 2012; Fattori et al., 2021). Upon activation, the peptidergic nociceptors subpopulation releases neuropeptides such as calcitonin gene-related peptide (CGRP) that signal through the calcitonin receptor-like receptor (CLR) in complex with receptor activity modifying protein 1 (RAMP1). The release of these neuropeptides by nociceptors shapes immune cell function in the innervated tissue in a context-dependent manner in a process known as neuroimmune communication (Baral et al., 2019; Chiu et al., 2012; Fattori et al., 2021). A better understanding of the consequences of neuropeptide release in different diseases is essential to determine the relevance of CGRP/RAMP1 targeting drugs for treating pain.

Endometriosis is a painful gynecological inflammatory disease that affects 10-15% of women (Parasar et al., 2017) and transgender men (Shim et al., 2020). The disease results in annual health care costs approaching $100 billion in the US alone (Soliman et al., 2018). Histologically, the defining characteristic of endometriosis is the ectopic presence of tissue (lesions) mimicking the appearance of uterine (eutopic) endometrium (Parasar et al., 2017). These lesions are defined by their endometrial-like glands, stroma, and hemosiderin with blood vessels, nerve fibers, muscle, and immune cells. Debilitating pain, including chronic pelvic pain, dysmenorrhea, dyspareunia, and dyschezia, are common clinical features of patients with endometriosis (Laux-Biehlmann et al., 2015; McKinnon et al., 2015). Pelvic pain represents a major clinical problem since women with that symptom report lower quality of life and mental health (Facchin et al., 2015; Soliman et al., 2018). Current treatments for pain in endometriosis are limited to non-steroidal anti-inflammatory drugs (NSAIDs), other analgesics, hormonal agents, and surgical removal of the lesions. While effective for a fraction of patients, hormonal therapies and NSAIDs can cause short and long-term side effects, and should be used with caution by patients with comorbidities (Soliman et al., 2018). This lack of treatment effectiveness contributes to endometriosis patients experiencing chronic opioid use, a diagnosis of opioid dependence/abuse, and opioid overdose (Chiuve et al., 2021). Therefore, new medical therapies and targets that provide long-term benefits are sorely needed.

A recent GWAS study shows a positive genetic correlation between migraines and endometriosis (rg=0.29, p=1.05x10^-16^) (Rahmioglu et al., 2023). This might suggest that endometriosis and migraine could share some pathophysiologic mechanisms. Human (Nishimoto-Kakiuchi et al., 2023; Tan et al., 2022) and mouse (Fattori et al., 2020a; Fusco et al., 2018; Ono et al., 2021) studies show that the lesion and peritoneal immune cell landscape is drastically changed during endometriosis, with macrophages being the main affected cells. In the peritoneal cavity (PerC), there are two main types of macrophages: an embryonic-derived type that is F4/80^hi^MHCII^lo^ and known as large peritoneal macrophages (LPM), and another that is bone-marrow-derived, F4/80^lo^MHCII^hi^, and known as small peritoneal macrophages (SPM, also CCR2^+^). LPM outnumbers SPM by ∼3-6-fold, a ratio that is changed toward SPM during abdominal/peritoneal inflammation due to an influx of inflammatory monocytes (Ghosn et al., 2010; Vega-Perez et al., 2021). However, the extent to which nociceptors mediate endometriosis disease outcome and monocyte recruitment is not known. Because endometriosis lesions are highly innervated, including the presence of CGRP^+^ fibers (Laux-Biehlmann et al., 2015; McKinnon et al., 2015), we hypothesized that neuroimmune communication is a driver of endometriosis-associated pain and lesion establishment/growth.

To test this hypothesis we used human samples and our validated mouse model of endometriosis (Fattori et al., 2020a). Mouse models of endometriosis-associated pain often require surgery to implant lesions, but abdominal surgery changes the phenotype of peritoneal cells, confounding analysis of endometriosis pathophysiology (Bain et al., 2020). Moreover, although spontaneous pain is the main complaint of patients with endometriosis, it is rarely reported/measured in endometriosis models (Simitsidellis et al., 2018). To address these issues, we used our validated non-surgical mouse model of endometriosis-associated pain (Fattori et al., 2020a), in which lesions are induced by injection and the resulting mice exhibit both evoked abdominal pain (von Frey) and spontaneous pain behaviors, including abdominal licking, contortions, and squashing (Fattori et al., 2020a).

We found that nociceptor to macrophage communication through CGRP/RAMP1 signaling supports endometriosis lesions and promotes pain. Furthermore, four different FDA approved drugs blocking CGRP/RAMP1 signaling reduced both evoked and spontaneous pain, as well as lesion size, Since the lack of drug efficacy at reducing ongoing pain drives most endometriosis therapy failure, these data showing efficacy with non-opioid and non-hormonal drugs might lead to clinical benefits for endometriosis patients.

## 2. Results

### 2.1 Human and mouse endometriosis lesions express CGRP^+^ fibers and endometriosis activates CGRP^+^ DRG neuron

The presence of CGRP-, TRPA1-, and TRPV1-expressing fibers and immune cells, such as macrophages, are commonly observed in endometriotic lesions (Bohonyi et al., 2017; Fattori et al., 2020a; Greaves et al., 2014; Mechsner et al., 2007; Tokushige et al., 2007). We have previously found that mouse endometriosis lesions contain TRPV1^+^, TRPA1^+^, and CGRP^+^ nociceptors (Fattori et al., 2020a). Using immunofluorescence, we confirmed that both mouse (Fig 1A) and human lesions contain CGRP^+^ fibers (Fig 1B). Furthermore, using mouse endometriosis lesion explants, we observed increased levels of soluble CGRP (Fig 1C). At the DRG level, we observed that CGRP^+^ neurons from endometriosis were more activated when compared to DRG neurons from sham mice (without lesions), as demonstrated by double staining for CGRP and phospho-NF-κB p65 (Fig 1D), a transcription factor that is activated during pain and inflammation. Our data, together with clinical evidence of a positive genetic correlation between migraine and endometriosis (Rahmioglu et al., 2023), indicates that CGRP is likely to play a role in endometriosis pathophysiology.

**Fig 1.**
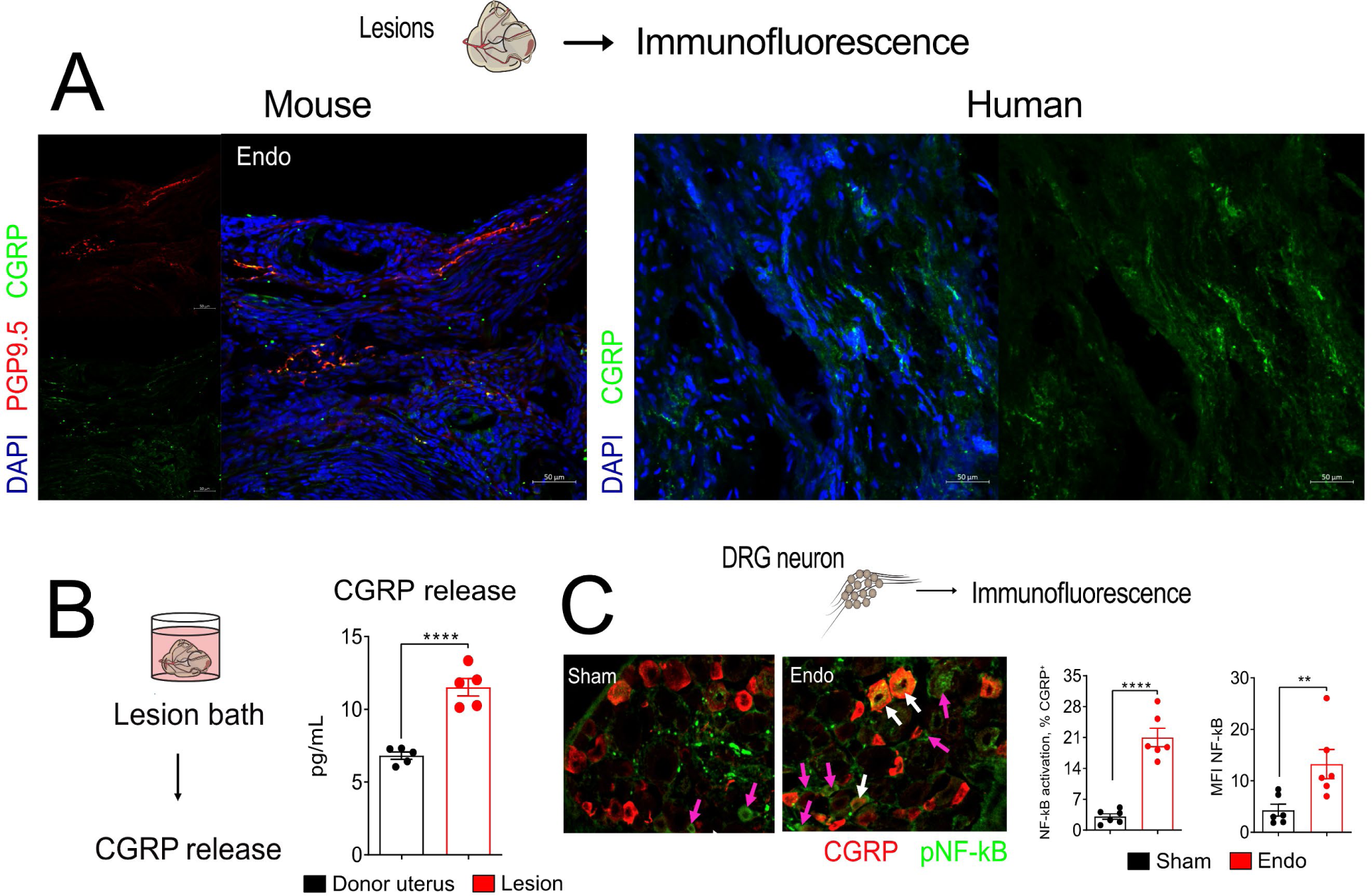
Human and mouse endometriosis lesions contain CGRP+ fibers and endometriosis activates CGRP^+^ DRG neurons. **(A)** Immunofluorescence staining of mouse and human endometriosis lesions for PGP9.5 (pan neuronal marker) and CGRP. **(B)** CGRP release in organ bath supernatants from endometriosis lesions collected at 56 dpi as measured by EIA. Results are expressed as mean ± SEM of 5 wells per group per experiment, two independent experiments (Student’s t-test, ****p<0.0001). **(C)** Immunofluorescence staining of phospho-NF-κB (p65 subunit) in CGRP+ DRG neurons from sham and endometriosis lesion bearing mice. Pink arrows point to pNF-κB^+^ DRG neurons. White arrows point to CGRP^+^pNF-κB^+^ DRG neurons. Results are expressed as mean ± SEM of 6 mice per group per experiment, two independent experiments (Student’s t-test, ****p<0.0001; **p<0.02).

### 2.2 Endometriosis lesions activate TRPA1^+^ and TRPV1^+^ nociceptors and induce CGRP release

Because nociceptors sense and respond to environmental cues and pathogen-derived products (Baral et al., 2019; Chiu et al., 2012; Fattori et al., 2021), we asked whether endometriosis lesions could directly activate nociceptors. Measurements of intracellular calcium in DRG neurons showed that lesion homogenate directly activate TRPA1^+^ (Fig 2A, a1) and TRPV1^+^ nociceptors (Fig 2A, a2). We then asked what molecules in the lesions could be responsible for this direct activation of nociceptors. A cytokine array revealed that 61 out the 111 mediators were differentially expressed in the lesions when compared to the donor uterus tissue used to induce endometriosis (Fig 3B, left). Similarly, an angiogenesis array revealed that 38 of 53 mediators were differentially expressed in the lesions when compared to the donor uterus (Fig 3B, right). In the arrays, we observed an increase in adhesion molecules such as P-selectin and VCAM-1, as well as an increase in CD14, a co-receptor for several toll-like receptors expressed by macrophages and other cells. Moreover, we found increased levels of the angiogenic factors vascular endothelial growth factor (VEGF), placental growth factor (PLGF), fibroblast growth factors (FGF), and platelet-derived growth factor (PDGF). We also found an increase in several macrophage regulators such as CCL2, CCL3, CX_3_CL1, and M-CSF (Fig. 2B). These data show that the lesion environment is very different than the donor uterus tissue. Due to VEGF and PLGF being implicated in the pathogenesis of endometriosis (Vodolazkaia et al., 2016) and their receptor (VEGFR1) being expressed by mouse DRG neurons (Micheli et al., 2021; Selvaraj et al., 2015), we next wondered whether VEGF and PLGF could contribute to the nociceptor activation we observed in our calcium imaging (Fig 2A). First, we confirmed that VEGFR1 is expressed by DRG neurons by immunohistochemistry (Fig 2C). Then we used a CGRP release assay in cultured DRG neurons to demonstrate that PLGF (Fig 2D, d.1) and VEGF (Fig 2D, d.2) increased soluble CGRP in a concentration-dependent manner, indicating that both molecules directly activate nociceptors and induce CGRP release.

**Fig 2.**
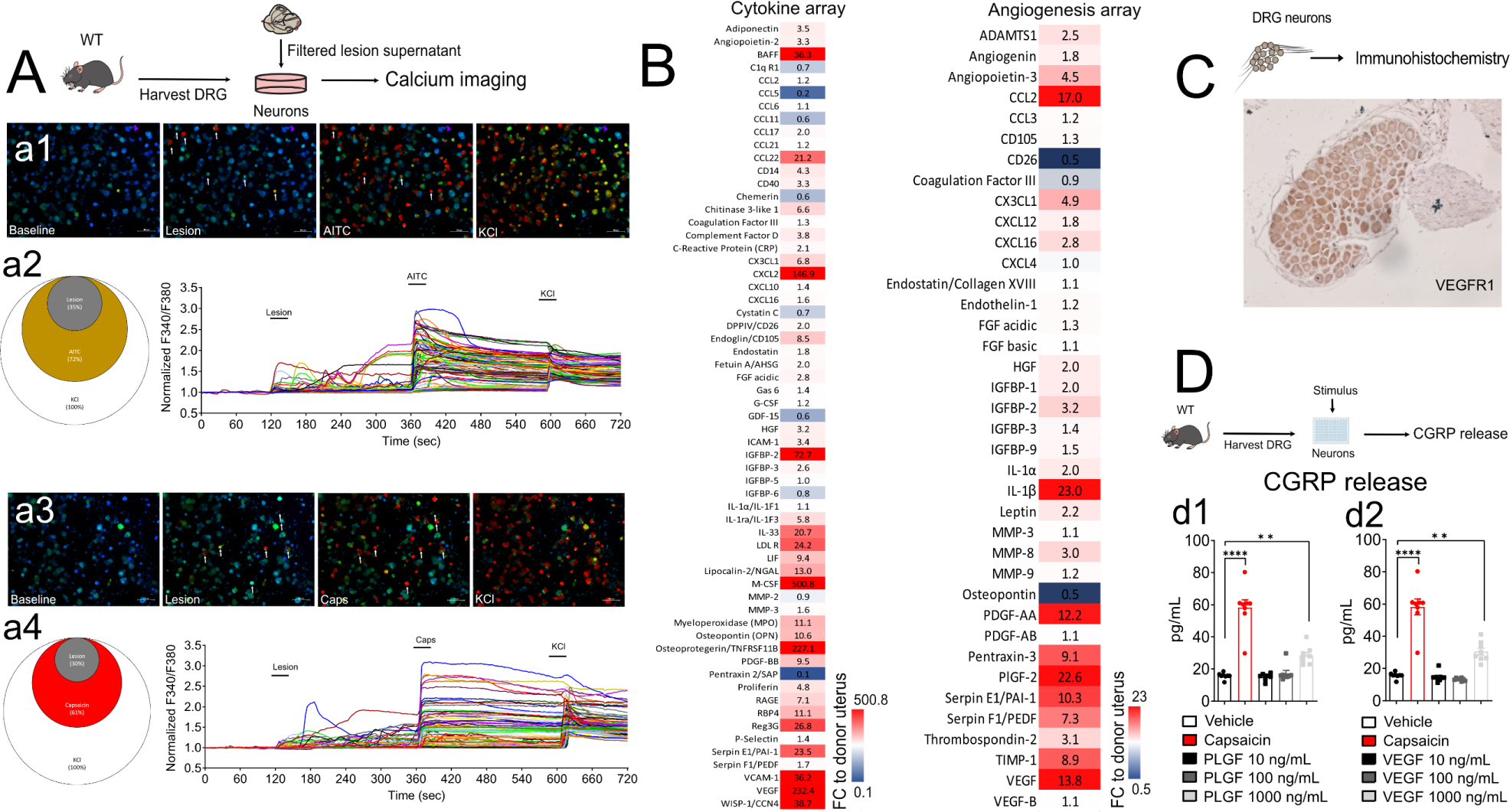
Endometriosis lesions activate TRPV1^+^ and TRPA1^+^ nociceptors and induce CGRP release. **(A)** Fura-2AM imaging of calcium influx into DRG neurons cultured from naive mice. Fluorescence ratio traces (panels **a2** and **a4**) represent calcium influx into DRG cells from representative fields (panel **a1** and **a3**) throughout the 12 min of recording. Results are expressed as mean fluorescence ratio representing intracellular calcium concentration at the baseline (zero-second mark) and following the stimulus with lesion supernatant (120s mark), TRPA1 agonist AITC **(a1, a2)** or the TRPV1 **(a3, a4)** agonist capsaicin (360s mark) and KCl (600 seconds mark, activates all neurons). The fraction of cell bodies responsive to each stimulus is represented by Venn diagrams (**a2** and **a4**). **(B)** Regulation of pro-inflammatory mediators as measured by cytokine and angiogenesis array kits. Endometriosis lesions were collected 56 dpi. Donor uterus was used as control. Results are expressed as fold-change (FC) vs. donor uterus. **(C)** VEGFR1 expression in DRG neurons from naive mice as determined by IHC and **(D)** measurement of CGRP release into the supernatant of cultured DRG neurons stimulated for 1 hour with PLGF **(d1)** or VEGF **(d2).** Concentrations were measured by EIA and results are expressed as mean ± SEM of 8 wells per group per experiment, two independent experiments (one-way ANOVA followed by Tukey’s post-test, ****p<0.0001); ***p<0.01).

**Fig 3.**
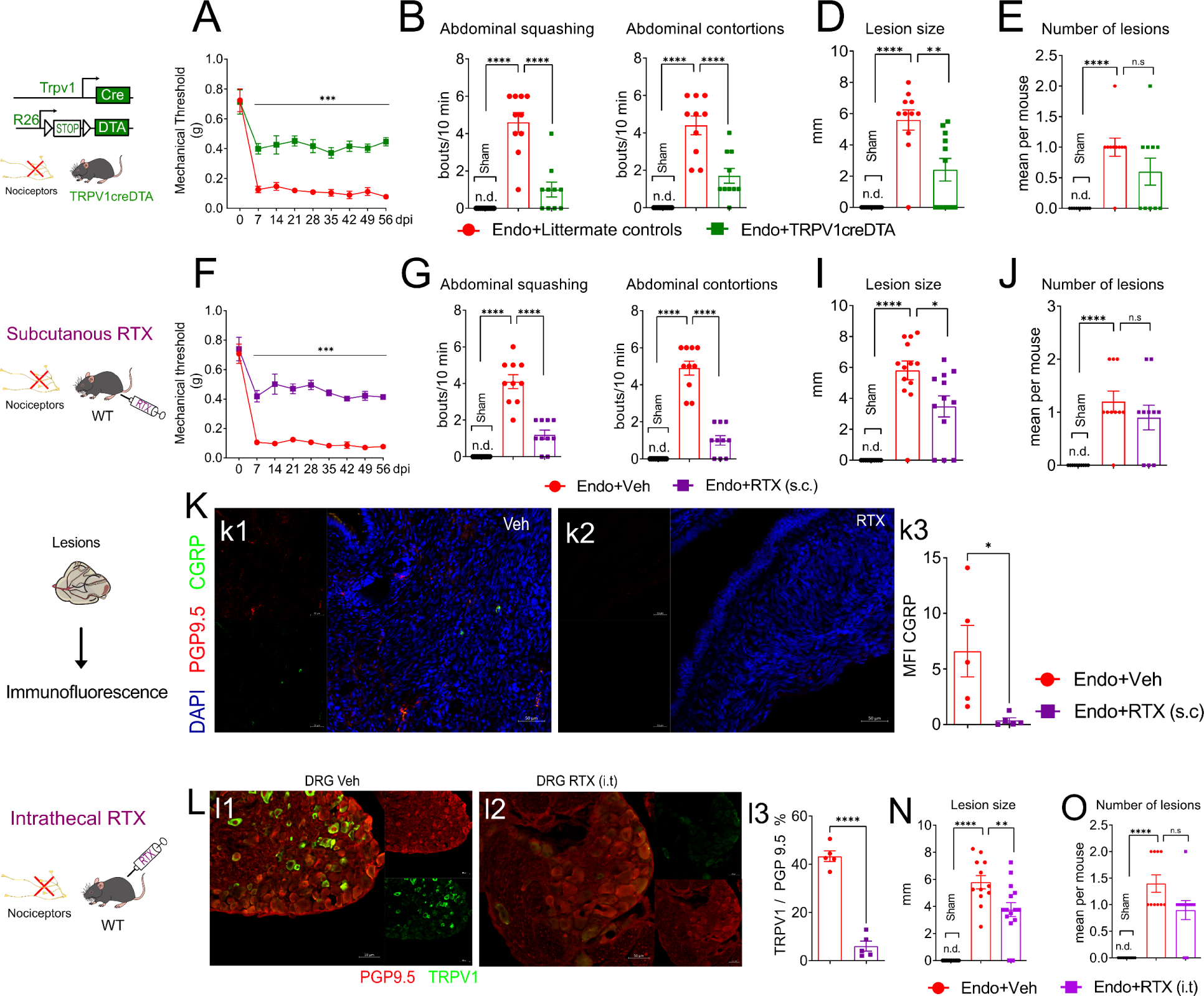
TRPV1^+^ nociceptors control endometriosis lesion establishment and growth. **(A and F)** Abdominal mechanical hyperalgesia measured using von Frey filaments before (zero) and after (7-56 dpi, weekly) endometriosis induction. Results are presented as mean ± SEM of mechanical threshold, n = 10 mice per group (two-way repeated-measures ANOVA followed by Tukey’s post hoc, ***p< 0.001). **(B and G)** Spontaneous pain measured by abdominal squashing and abdominal contortions. For abdominal squashing, the number of times the mice pressed the lower abdominal region against the floor was quantified for 10 minutes. Sham mice do not display abdominal squashing. Abdominal contortions were quantified for 10 minutes by counting the number of contractions of the abdominal muscle together with stretching of hind limbs. Sham mice did not display abdominal contortions. **(D, I, and N)** Lesion size, as determined in the remaining lesions by measuring perpendicular diameters. **(E, J, and O)** Number of visible lesions was calculated as sum of total lesions per mouse. Sham mice do not show any lesion (n.d. = not detectable). Results are expressed as mean ± SEM, n = 10 (one-way ANOVA followed by Tukey’s post hoc, ****p< 0.0001). **(K)** The effect of subcutaneous RTX treatment on the number of CGRP fibers in lesions 28 dpi as measured by immunofluorescence staining. Representative pictures are display in panels **k1** and **k2** while panel **k3** show the quantification of CGRP staining upon RTX ablation (Student’s t-test, *p<0.05). **(L)** The effect of intrathecal RTX induced nociceptor ablation on lesion size. Comparing **l1** and **l2** immunofluorescence stained for PGP9.5 and TRPV1 confirms nociceptor ablation (one-way ANOVA followed by Tukey’s post hoc, ****p<0.0001, **p<0.02). MFI: mean fluorescence intensity.

### 2.3 TRPV1^+^ nociceptor signaling control endometriosis lesions establishment and growth

We then asked whether nociceptors play a role in endometriosis. For that, we bred TRPV1cre with Rosa26 *lox*-STOP-*lox* DTA mice (ROSA-DTA) to create TRPV1creDTA nociceptor ablated mice and Cre^−^ littermate (LM) controls (Fig 3A). We found that targeted ablation of TRPV1^+^ nociceptors using TRPV1creDTA mice reduced mechanical threshold (Fig 3A), abdominal squashing and contortions (Fig 3B). Surprisingly, these mice also showed reduced lesion implantation (Fig 3C) and significantly smaller lesions when compared to the littermate controls (Fig 3D), indicating that nociceptors control not only pain, but also lesion size.

To confirm that this is caused by the absence of TRPV1-expressing cells, rather than e.g., developmental feedback, we next performed chemical ablation of nociceptors with resiniferatoxin (RTX). RTX is a high-affinity agonist for TRPV1 that causes acute chemical ablation of nociceptors (Baral et al., 2018; Chiu et al., 2013). Targeted TRPV1 ablation was confirmed by immunofluorescence of the lesions (Fig 3K) and DRG neurons (Fig. 3L). We found that systemic (subcutaneous) RTX treatment reduced evoked mechanical pain (Fig 3F) and spontaneous pain (Fig. 3G). Similar to the TRPV1creDTA mice, we found that RTX treatment also reduced lesion size (Fig. 3I and N), indicating that TRPV1^+^ nociceptors are important for lesion establishment and growth. To determine that nociceptors with cell bodies in the DRG (rather than, e.g., nodose ganglia) are responsible for the lesion growth, we next used intrathecal injection of RTX and found similar results (Fig 3L-O). This indicates that TRPV1^+^ nociceptors control endometriosis lesion growth and establishment.

We next asked whether action potential propagation is required for the lesion-supporting effect of nociceptors. To that end, we tested QX-314, a permanently charged quaternary derivative of lidocaine (Fig 4). QX-314 blocks nociceptor activation and CGRP release only when large pore channels, such as TRPV1, are activated (Binshtok et al., 2007; Talbot et al., 2015). Consistent with our TRPV1^+^ nociceptor ablation results, we found that inhibiting nociceptor activation with QX-314 reduced mechanical threshold (Fig 4A) and spontaneous pain in a dose-dependent manner (Fig 4B). Importantly, we also found that QX-314-treated mice had significantly smaller lesions (Fig 4D) and fewer lesions when compared to vehicle-treated animals (Fig 4E), confirming that TRPV1^+^ neuron signaling supports not only the pain caused by lesions, but also lesions themselves.

**Fig 4.**
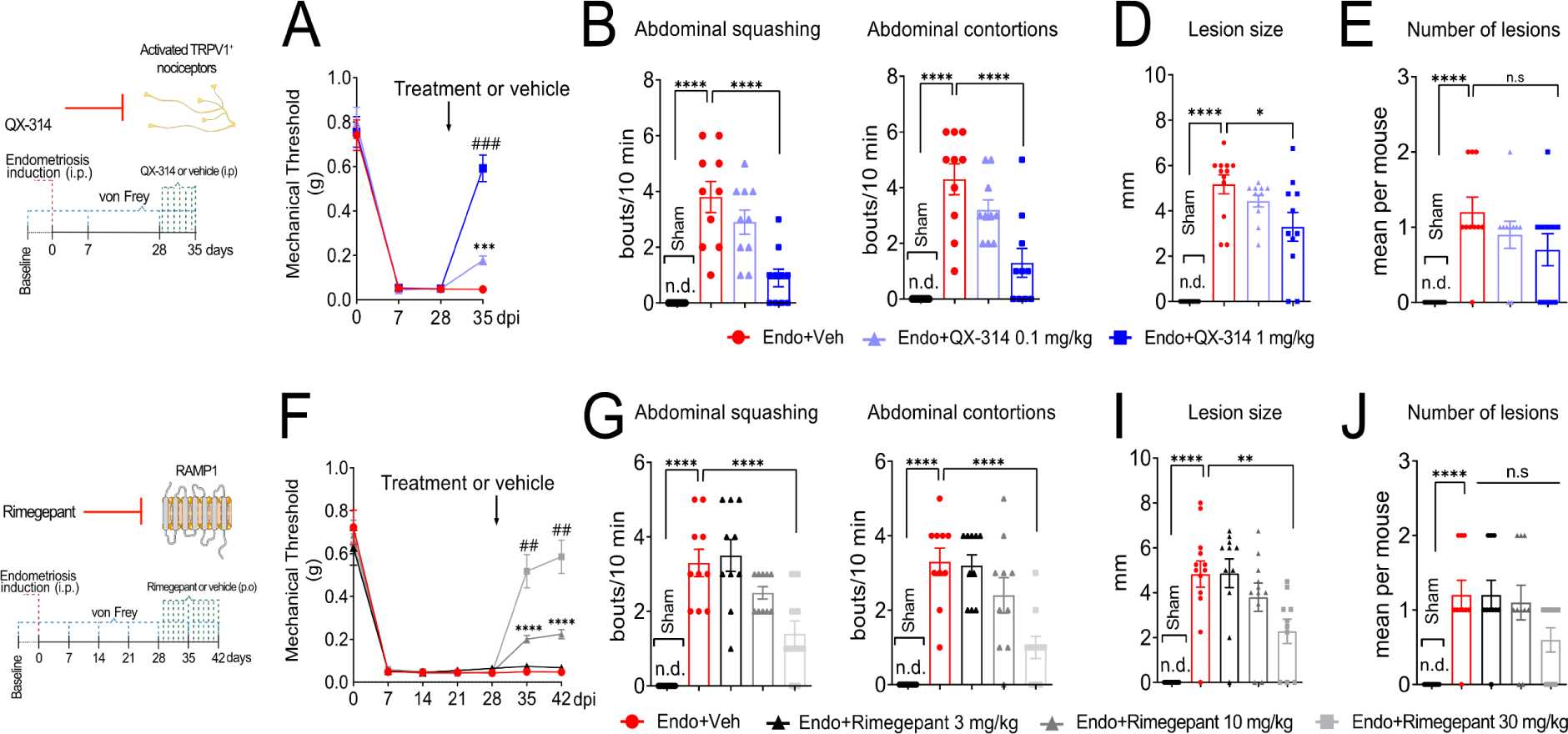
Blocking nociceptor activation and CGRP/RAMP1 signaling reduces endometriosis-associated pain and lesion size. **(A and F)** Abdominal mechanical hyperalgesia measured before (zero) and after (weekly) lesion induction using von Frey filaments. Results are presented as mean ± SEM of mechanical threshold, n = 10 mice per group (two-way repeated-measures ANOVA followed by Tukey’s post hoc, ****p< 0.0001; ***p=0.002; ###p = 0.0001 vs QX-314 0.1 mg/kg; ### p< 0.0005 vs Rimegepant 10 mg/kg). **(B and G)** Spontaneous pain measured by abdominal squashing and abdominal contortions as described in Figure 3 B and G. **(D and I)** Lesion size as determined by measuring perpendicular diameters. **(E and J)** Number of visible lesions was measured as sum of total lesions per mouse. Sham mice do not show any lesion (n.d. = not detectable). Results are expressed as mean ± SEM, n = 10 (one-way ANOVA followed by Tukey’s post hoc, ****p< 0.0001; **p<0.02; *p<0.05).

### 2.4 Blocking CGRP/RAMP1 signaling reduces endometriosis-associated pain and lesion size

Because CGRP is a key mediator of neuro-immune communications that is released by activated nociceptors, we next asked whether blocking CGRP/RAMP1 signaling with drugs that either prevent CGRP release or prevent CGRP from binding to its coreceptor, RAMP1, would affect endometriosis pain and lesion growth. We used 4 different FDA approved drugs that target either CGRP (fremanezumab or galcanezumab – anti-CGRP monoclonal antibodies) or RAMP1 (rimegepant or ubrogepant – RAMP1 antagonist small molecules). We observed that treatment with rimegepant reduced mechanical threshold (Fig 4F) and spontaneous pain (Fig 4G) in a dose-dependent manner. Importantly, rimegepant also reduced lesion size (Fig 4I). Similarly, ubrogepant, fremanezumab, and galcanezumab all reduced mechanical threshold (Fig S1A), and spontaneous pain (Fig S2B). At the chosen dose ubrogepant also reduced lesion size (Fig S1D). This suggests that this class of drugs, which are currently approved to treat migraines, will be effective at treating endometriosis pain as well as at reducing lesion size.

In line with our cytokine and angiogenesis array data, we confirmed by ELISA that CCL2, CCL3, CX_3_CL1, as well as PLGF and VEGF, are increased in both the lesions (Fig 5A) and the PerC wash (Fig 5B) of endometriosis lesion-bearing mice when compared to lesion-free mice (sham or donor uterus used as controls). Importantly, rimegepant reduced the levels of CCL2, CLL3, PLGF, and VEGF in the lesions (Fig 5A), while also reducing levels of CCL3 and CX_3_CL1 in the PerC wash (Fig 5B). This indicates that blocking CGRP/RAMP1 signaling not only improves endometriosis pain, but also changes lesion microenvironment and size.

**Fig 5.**
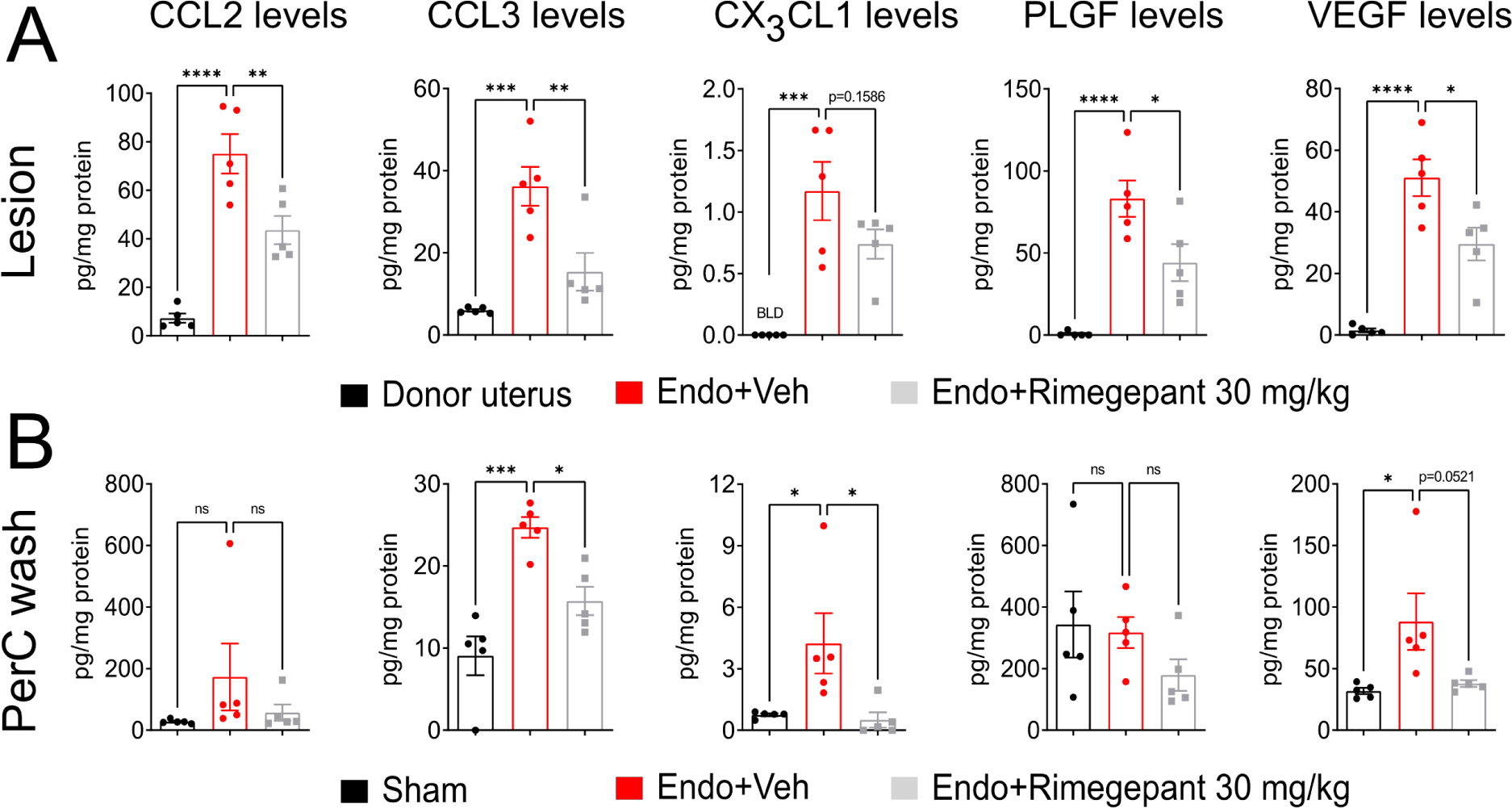
Rimegepant reduces endometriosis-induced chemokines and growth factor increases in the lesion and peritoneal cavity wash. **(A)** ELISA-measured concentrations of CCL2, CCL3, CX3CL1, PLGF, and VEGF in the lesions of vehicle- and rimegepant (30 mg/kg)-treated mice. Donor uterus (tissue used to induce endometriosis) was used as control since sham mice do not develop lesions. **(B)** ELISA-measured concentrations of CCL2, CCL3, CX3CL1, PLGF, and VEGF in the peritoneal wash of sham-injected mice, and lesion-bearing mice treated with vehicle or rimegepant (30 mg/kg). Results are expressed as mean ± SEM, n = 10 (one-way ANOVA followed by Tukey’s post hoc, ****p< 0.0001; **p<0.02; *p<0.05).

### 2.5 TRPV1^+^ nociceptors control monocyte recruitment during endometriosis and facilitate lesion growth

Having observed that CGRP blockers reduced lesions and pain and mindful of the demonstrated role of the immune/inflammatory microenvironment in endometriosis, we hypothesized that nociceptors may directly signal to immune cells via neuropeptides to facilitate lesion establishment/growth and pain. To test this hypothesis, we started by performing single-cell RNA sequencing (scRNAseq) analysis from the PerC wash of TRPV1creDTA mice, sham littermates (lesion free mice), and lesion bearing littermate controls (Fig 6A, 7A, and S2-9). After defining cluster identities by comparing transcriptional markers with published PerC single-cell datasets (ImmGen, 2016; Lantz et al., 2020; Liu et al., 2021), we observed a repertoire of myeloid and lymphoid cells (Fig 6A and S2-9). Nine different clusters were identified with 4 of them being different types of macrophages, including LPM (F4/80^hi^MHCII^lo^, and *Gata6*^hi^ and *Timd4*^hi^) and SPM (F4/80^lo^MHCII^hi^ and positive for *Ccr2*^+^) (Fig 6A and S2-9). Differentially expressed genes (DEGs) in the different macrophage clusters represent ∼68% of total DEGs (Fig S2B and C), indicating that macrophages are the main cells affected during endometriosis. While SPM and LPM clusters represent ∼40% of total DEGs, we did not find any DEGs when comparing sham to endo (lesion-bearing) mice in the proliferating macrophage and neutrophil clusters (Fig S2B and S8). This indicates that SPM and LPM play a major role in the disease while proliferating macrophages and neutrophils contribute little to the aberrant disease microenvironment. Gene Ontology (GO) enrichment analysis highlighted differential regulation of genes involved in the biological processes such as leukocyte chemotaxis, migration, and mononuclear cell proliferation (Fig 6B), thus further highlighting the roles of these cells in endometriosis lesion formation. We specifically found 85 upregulated DEGs in the LPM cluster when compared to sham, including chemokines such as *Ccl6*, *Ccl9*, and *Arg1* (Fig. 6C left and S2B). Similarly, we found 62 upregulated genes in the SPM cluster including the cytokine *Il1b*, genes related to cell migration *S1006a*, and a *Cebpb* (Fig 6C right and S2B), which is a promoter implicated in the polarization into a subset of endometriosis-permissive macrophages (Ruffell et al., 2009). Since a shift in LPM/SPM ratio is often associated with PerC inflammation (Ghosn et al., 2010; Vega-Perez et al., 2021), we then wanted to determine whether this would happen in our mouse model. We found that lesion-free sham and lesion-bearing TRPV1creDTA mice had similar percentages of SPM, while lesion-bearing LM control mice had increased the percentage of SPM (Fig. 7B). Specifically, LPM/SPM ratio for sham mice was 6.40%, while for LM control was 3.32%, and for TRPV1creDTA mice was 4.95% (Fig 7B). This demonstrates that endometriosis changes the LPM/SPM ratio and targeted ablation of TRPV1^+^ nociceptors rescues this change, suggesting that nociceptors mediate SPM recruitment to the PerC.

**Fig. 6.**
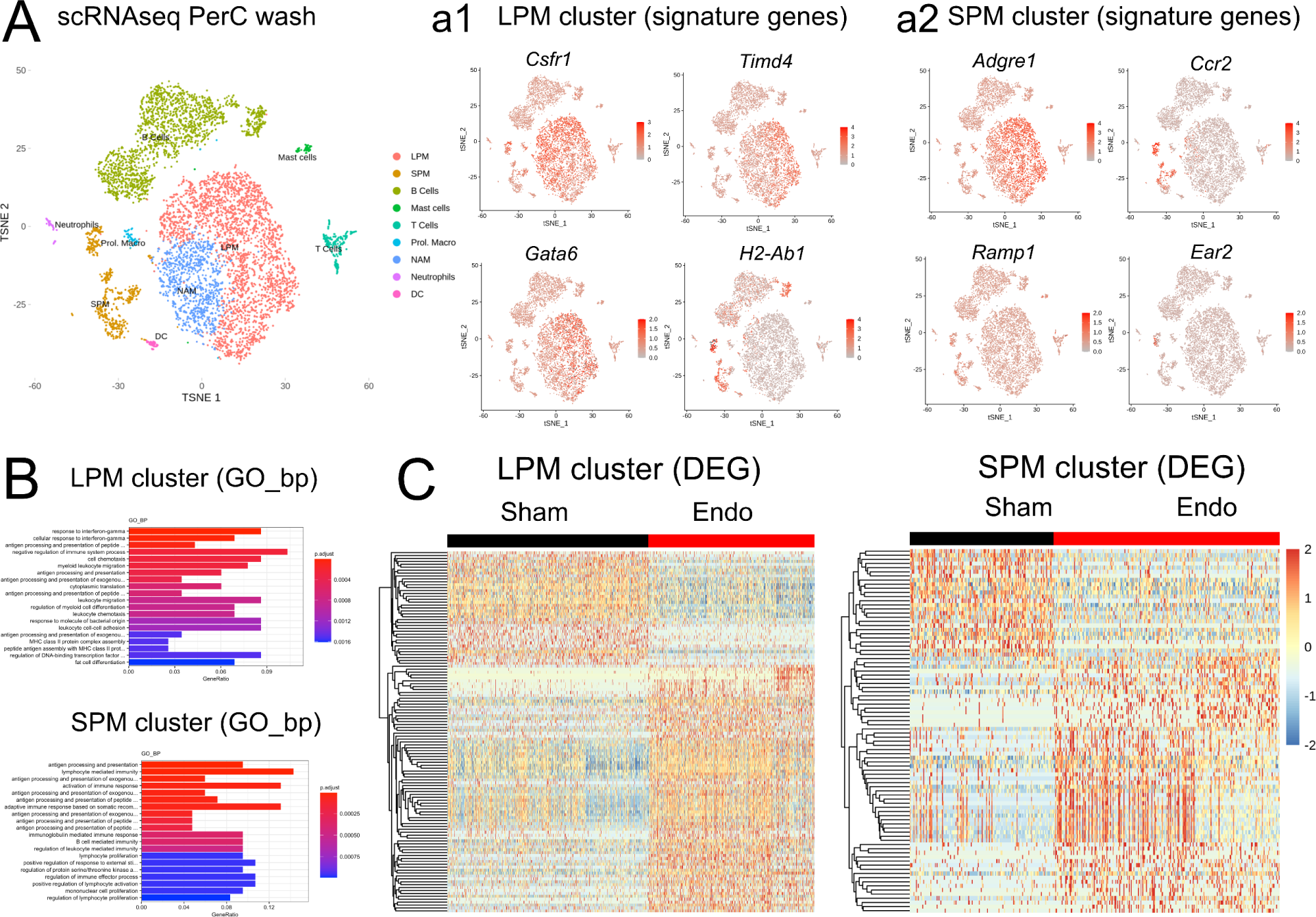
RNAseq analysis of PerC immune cells with and without endometriosis lesions. PerC wash of sham and lesion-bearing mice were collected at 28 dpi for scRNAseq analysis. **(A)** tSNE plot displaying the nine different populations of PerC immune cells. tSNE subplots displaying signature genes for the **(a1)** LPM and **(a2)** SPM clusters. **(B)** GO enrichment analysis of the LPM (top) and SPM (bottom) clusters in sham vs endometriosis mice. **(C)** Heatmap showing DEGs of the LPM (left) and SPM (right) clusters in sham vs endometriosis mice. (n = 5 pooled mouse PerC washes for each condition).

**Fig 7.**
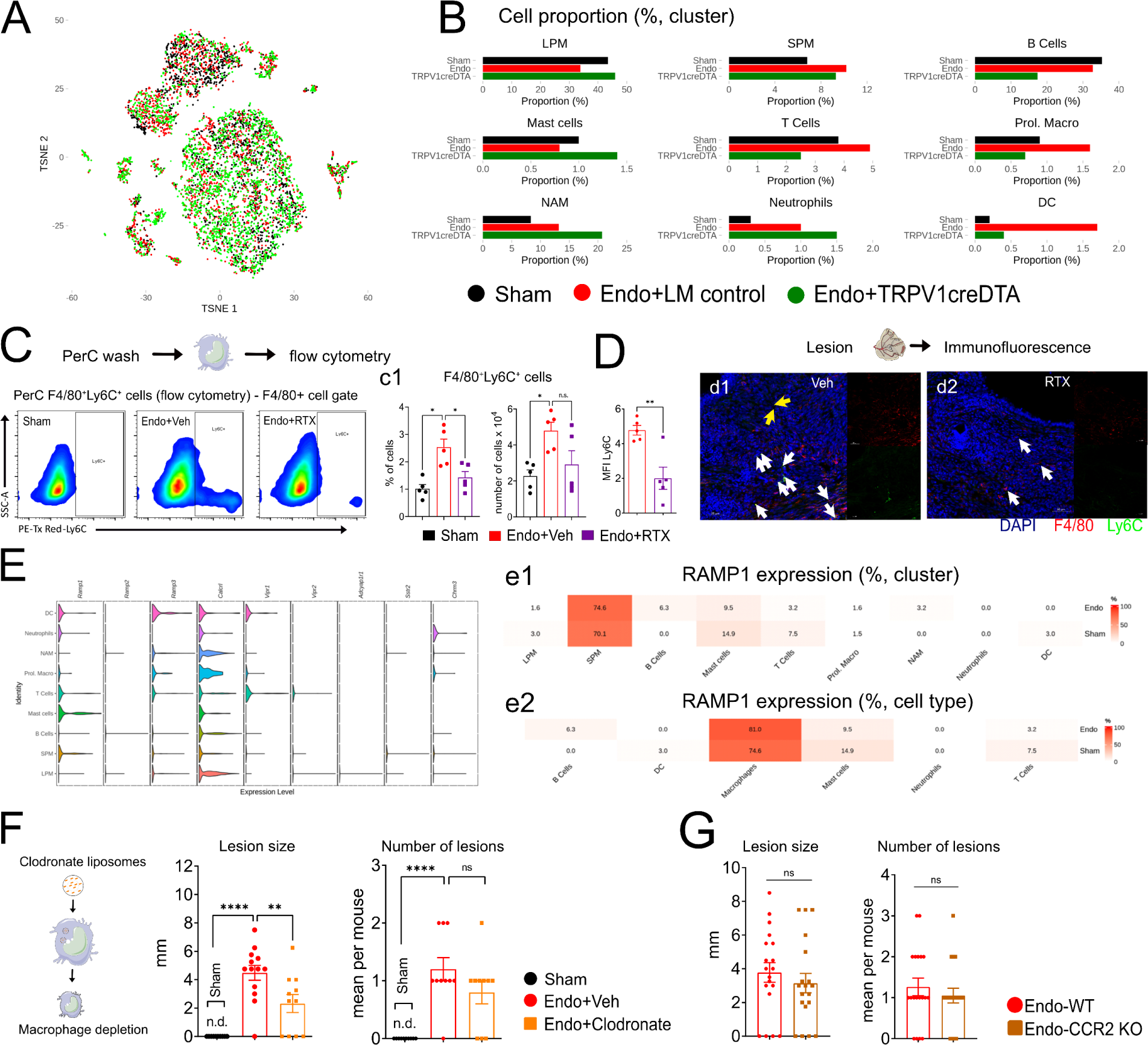
Nociceptors control monocyte recruitment to the peritoneal cavity and lesions during endometriosis. **(A, B, and E)** PerC wash of sham littermate (healthy), littermate lesion-bearing control, and lesion-bearing TRPV1-creDTA mice were collected at 28 dpi for scRNAseq analysis. **(A)** tSNE plot of PerC wash cells collected from wash of sham (steady state), LM control, and TRPV1-creDTA mice. **(B)** Cell proportion (percentage for each cluster) within the three groups. **(C)** Flow cytometry measurement of monocyte recruitment to the PerC of sham, vehicle-, or RTX-treated mice 28 dpi. **(D)** Immunofluorescence staining of recruited monocytes in lesions from the same mice. White arrows show F4/80^+^Ly6C^+^ cells while yellow arrows Ly6C^+^ only cells Results are expressed as mean ± SEM, n = 5 (one-way ANOVA followed by Tukey’s post hoc, **p<0.02, *p<0.05). **(E)** Violin plots showing the gene expression levels per cluster and the percentage of cells from each cluster expressing genes for neuropeptide receptors. Panel **e1** shows the percentage that each cell cluster and **e2** displays the percentage that each cell type represents among the RAMP1^+^ cells. **(F)** The impact of clodronate-induced macrophage depletion on endometriosis lesion size. **(G)** Lack of effect of CCR2 knockout on lesion size or number. Lesion size was determined in the remaining lesions by measuring height and weight. **(F and G, left)** Lesion size was determined in the remaining lesions by measuring height and weight. **(F and G, right)** Number of visible lesions was calculated as sum of total lesions per mouse. Sham mice do not show any lesion (n.d. = not detectable). Results are expressed as mean ± SEM, n = 10 (F) or 19 (G) (one-way ANOVA followed by Tukey’s post hoc, ****p< 0.0001; **p<0.02; *p<0.05). MFI: mean fluorescence intensity.

To determine whether targeted ablation of nociceptors would indeed result in less monocyte recruitment, we next performed flow cytometry from the PerC wash of sham, vehicle-treated, and RTX-treated mice. We found an increased percentage of F4/80^+^Ly6C^+^ cells in the PerC wash of vehicle-treated mice; RTX treatment normalized this percentage (Fig 7C). To extend these results to lesions themselves, we performed immunofluorescence on lesions from vehicle- and RTX-treated mice. We found fewer F4/80^+^Ly6C^+^ cells in the nociceptor ablated mice (Fig 7D). These data confirm that TRPV1^+^ nociceptors control monocyte recruitment to the PerC during endometriosis. Given that targeted ablation of nociceptors reduced monocyte recruitment, we then surveyed in our cell clusters for the expression of neuropeptide receptors (Fig. 7E). We found that *Ramp1* and *Calcrl* are expressed by most clusters, with the SPM cluster accounting for approximately 75% of the RAMP1^+^ cells (Fig 7E and e1, top heatmap). When combining all four clusters, macrophages account for 81% of the RAMP1^+^ cells (Fig 7E and e1, bottom heatmap). Based on these data, we propose that macrophage recruitment is a key consequence of nociceptor activity.

To measure the role of macrophages in lesion size, we depleted PerC macrophages using clodronate liposomes (Fig 7F). Intraperitoneal injection of clodronate liposomes significantly reduced the number of macrophages as demonstrated by a dramatic decrease in the number of CD45^+^F4/80^+^ cells in the PerC wash (Fig. S9A and B). We next induced lesions in vehicle- and clodronate-treated mice and found that the macrophage-depleted mice had smaller lesions (Fig. 7F), which indicates that macrophages are important for lesion growth and/or establishment. In addition to neuropeptides, nociceptors also release chemokines, such as CCL2, that drive monocyte recruitment to the inflammatory foci (Baral et al., 2019; Chiu, 2017). Because we observed that targeted ablation of nociceptors leads to a reduction in monocyte recruitment and the SPMs are the main RAMP1-expressing cells, we next sought to determine whether other nociceptor-derived molecules could recruit monocytes to the PerC. Therefore, we induced endometriosis lesions in CCR2 knockout mice to determine whether the absence of CCR2/CCL2 signaling would impair monocyte recruitment. Surprisingly, we found that lesion implantation and lesion size (Fig. 7G) were the same in knockout (KO) vs wild-type (WT) mice. Since CX_3_CL1 also drives monocyte recruitment, we next sought to determine whether CX_3_CL1/CX_3_CR1 signaling could be driving monocyte recruitment in our model. We induced endometriosis in CX_3_CR1^GFP^ reporter mice, a mouse strain in which a green fluorescent protein (GFP) sequence replaces the CX_3_CR1 gene, knocking out CX_3_CR1. Similar to our CCR2-KO data, we found no difference in lesion size or number of mice with lesions in the CX_3_CR1^GFP^ mice (Fig. S9C). Given the continued macrophage recruitment in the absence of chemokine signaling and reduction of that recruitment upon nociceptor ablation, our data suggest that nociceptors dictate macrophage recruitment via CGRP release for pain and lesion formation during endometriosis via a process that is independent of CX_3_CR1 and CCR2 signaling.

### 2.6 CGRP/RAMP1 signaling on macrophages supports endometriosis lesion growth

Based on our scRNAseq data, we hypothesize that CGRP/RAMP1 signaling in macrophages is key for endometriosis pain and lesion formation. By immunofluorescence, we found that PerC macrophages from mice with endometriosis showed increased staining for pCREB, a transcription factor downstream RAMP1 signaling (Fig 8A, a1). We also confirmed by immunofluorescence the presence of RAMP1 both in mouse (Fig 6A, a3) and human (Fig 8A, a3) endometriosis lesions. We found that RAMP1 was mainly colocalized with F4/80^+^ cells, indicating that RAMP1 is mainly expressed by macrophages (Fig. 8A, a2). Since we observed that the lack of CCR2 does not interfere with lesion establishment/growth and that the SPMs (CCR2^+^ cells) account for most of the RAMP1^+^ cells in the PerC, we then wanted to confirm the presence of these cells in the lesions. To do that, we induced endometriosis in CCR2^RFP^ reporter mice, a mouse strain which a red fluorescent protein (RFP) sequence replaces the CCR2 gene. Using immunofluorescence, we found an increased number and percentage of RFP^+^ cells in the PerC of mice with endometriosis when compared to sham mice (Fig 8B, b1 and b2). In the lesions, we observed that these CCR2^RFP^ macrophages were in close proximity to CGRP^+^ fibers (Fig 8B, b3), indicating that these cells might be responsive to CGRP and CCR2 is not essential for their recruitment. To gain insights on how CGRP affects macrophages’ response to lesions, we cultured PerC macrophages with CGRP as a stimulus and performed an efferocytosis assay using apoptotic bodies from an immortalized endometriosis epithelial cell line (12Z) tagged with GFP. We observed that CGRP-treated macrophages had reduced efferocytosis ability when compared to vehicle-treated macrophages (Fig. 8C), indicating that CGRP reduces the capacity of macrophages to clear lesion debris.

**Fig 8.**
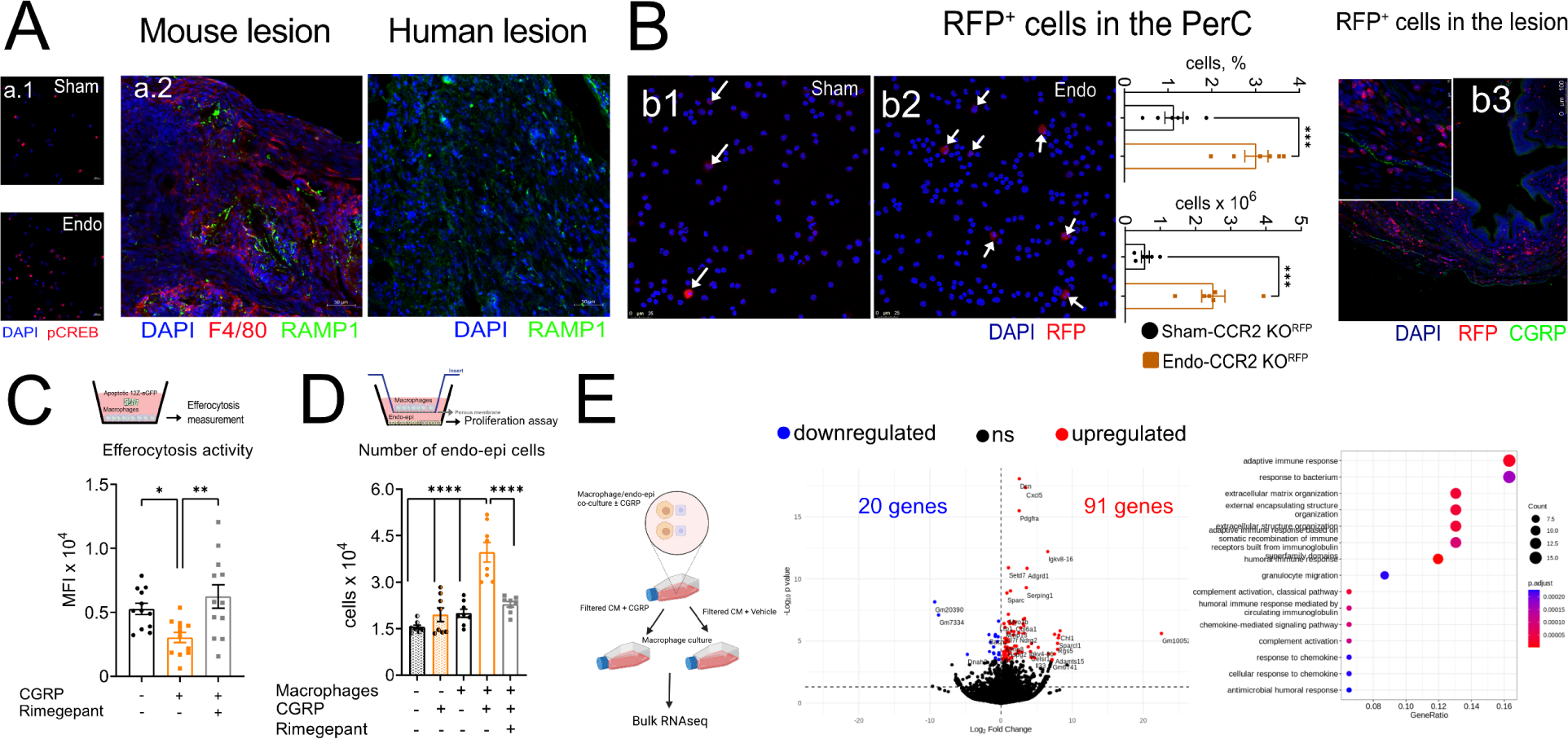
CGRP/RAMP1 signaling in macrophages supports endometriosis lesion growth. **(A)** Immunofluorescence staining for indicators of CGRP signaling. **(a1)** PerC wash from sham or endometriosis mice stained for pCREB to identify cells with active GPCR (i.e., RAMP1) signaling as determined by the presence of phospho-CREB. **(a2)** Mouse and human lesions stained for RAMP1 expression. **(B)** The presence of RFP-labeled macrophages in 28 dpi lesions of CCR2^RFP^ mice **(b1** and **b2**, white arrows) in PerC wash from sham and endometriosis mice and **(b3)** lesions from endometriosis mice. Results are expressed as mean ± SEM, n = 6 (one-way ANOVA followed by Tukey’s post hoc, ****p< 0.0001; **p<0.02; *p<0.05). **(C)** CGRP (100 nM for 24 hours) inhibits macrophage efferocytosis of etoposide-induced 12Z-eGFP apoptotic bodies in a RAMP1-dependent manner. Cells were washed 3 times with DPBS to remove non-engulfed 12Z-eGFP and then and fluorescence intensity determined. Treatment with rimegepant (100 nM) or vehicle was performed 30 min before stimulus with CGRP. **(D)** The effect of 100 nM CGRP-treated macrophages on endo-epi cell proliferation over 24h, measured using CyQuant. Treatment with Rimegepant (100 nM) or vehicle was performed 30 min before stimulus with CGRP. **(E)** Bulk RNAseq of the stimulated PerC macrophages with conditioned media (CM) from the vehicle- or CGRP-stimulated co-cultured cells (n = 2 for each condition). Results are expressed as mean ± SEM, n = 5-12 (one-way ANOVA followed by Tukey’s post hoc or Student’s t-test, ****P<0.0001, p<0.001, **p<0.02, *p<0.05). MFI: mean fluorescence intensity.

We next asked how macrophages might affect lesion expansion. We performed co-culture experiments using PerC macrophages and a mouse endometrial epithelial cell line (endo-epi cells). We found that CGRP-treated macrophages stimulated the growth of endo-epi cells (Fig. 8D), indicating that not only did CGRP impair the ability of macrophages to engage in efferocytosis, but also induced the release of factors that support endometrial cell growth. Notably, the reduction in efferocytosis (Fig 8C) and increase in endometrial cell growth (Fig 8D) was rescued by rimegepant, indicating that it is RAMP1-dependent. To identify soluble mediators that could be driving endometrial cell growth, we co-cultured macrophages and endo-epi cells with either CGRP or vehicle as a stimulus, and then performed bulk RNAseq of PerC macrophages stimulated with conditioned media (CM) from the vehicle- or CGRP-stimulated co-cultured cells (Fig. 8E). We found a total of 111 DEGs, 91 upregulated and 20 downregulated by CGRP (Fig 8E). The chemokines *Ccl2*, *Ccl7*, *Cxcl3*, *Cxcl5*, and *Cxcl13* as well as the cytokine *Il33* were upregulated by CGRP. Furthermore, we found an increase in growth factor binding proteins such as *Igfbp3* (insulin-like growth factor binding protein 3), *Igfbp4,* and *Igfbp7,* as well as *Pdgfra* (platelet-derived growth factor receptor A). Moreover, CGRP-treated macrophages showed increased expression of *Mmp3* (matrix metalloproteinases 3), *Mmp8*, *Mmp12*, and *Mmp13*. Several of these transcripts matched the data from our cytokine and angiogenesis array, including IGFBP3, CCL2, IL-33, MMP3, and MMP8 (Fig 2B). Interestingly, we also found that CGRP upregulated receptors to mediators whose levels were increased in our arrays such as *Il1r1* and *Pdgfra* (*i.e.* IL-1β and PDGF, respectively). Therefore, while not produced by macrophages during endometriosis, CGRP did increase macrophage responsiveness to these molecules. Altogether, indicating that CGRP changes the phenotype of macrophages to a “pro-endometriosis” state.

## Discussion

Lesion and peritoneal immune cell landscapes are drastically changed during endometriosis (Fattori et al., 2020a; Nishimoto-Kakiuchi et al., 2023; Ono et al., 2021; Tan et al., 2022). However, the extent to which the control of immune cells by nociceptors plays a role during endometriosis and whether targeting this communication effectively treats endometriosis is not yet known. Here we describe a nociceptor to macrophage communication pathway that drives endometriosis lesion growth and pain. We show that CGRP programs macrophages to a pro-endometriosis phenotype. CGRP impairs the efferocytosis ability of macrophages and induces macrophages to promote endometrial cell growth stimulation. At the transcriptional level, CGRP upregulates chemokines, cytokines, and enzymes that can support endometriosis lesion formation and pain. Importantly, rimegepant (FDA approved RAMP1 antagonist) rescues CGRP deleterious effects by normalizing the efferocytosis capacity and reducing endometrial cell growth. Moreover, blocking CGRP/RAMP1 signaling with 4 different FDA-approved drugs reduces pain (evoked and spontaneous) and lesion size. We also found that rimegepant normalizes chemokine and growth factors production in the lesions and PerC wash. Therefore, blocking neuroimmune communication by disrupting CGRP/RAMP1 signaling should be explored as a non-hormonal and non-opioid treatment for endometriosis.

Immune cell recruitment and function is significantly affected by tissue innervating nociceptors (Baral et al., 2019; Chiu et al., 2012; Fattori et al., 2021). Nociceptor-derived CGRP acting on macrophages (Chiu et al., 2013; Zaninelli et al., 2022), γδT cells (Baral et al., 2018), neutrophils (Baral et al., 2018; Pinho-Ribeiro et al., 2018), and DCs (Hanc et al., 2023; Riol-Blanco et al., 2014) worsens outcome in a variety of diseases. Here we show that during endometriosis nociceptors control macrophage function and recruitment to the PerC and lesions. In the PerC, there are two main types of macrophages: one embryonic derived that is F4/80^hi^MHCII^lo^ known as large peritoneal macrophages (LPM) and the other that is bone marrow derived and F4/80^lo^MHCII^hi^, known as small peritoneal macrophages (SPM) (Cassado Ados et al., 2015). In healthy animals, LPMs outnumber SPMs, but the SPM/LPM ration increases during peritoneal inflammation (Cassado Ados et al., 2015). In endometriosis, lesion macrophages are tolerogenic, pro-angiogenic, and pro-neurogenic (Tan et al., 2022). In fact, ablation of macrophages in different mouse models of endometriosis also resulted in smaller lesions (Bacci et al., 2009; Ono et al., 2021). We found that RAMP1 is mainly expressed by the CCR2^+^ macrophages (SPM cluster), indicating that this could be a nociceptor-responsive population. Corroborating this idea, targeted ablation of nociceptors reduced F4/80^+^Ly6C^+^ monocyte recruitment to the PerC and endometriosis lesions. Nociceptors release chemokines that also drive monocyte recruitment (Baral et al., 2019; Chiu et al., 2012). To determine the extent to which other nociceptor-derived mediators dictate monocyte recruitment to the PerC, we then induced endometriosis in CCR2-KO and CX_3_CR1-KO mice. Surprisingly, a lack of CCR2 or CX_3_CR1 did not affect macrophage recruitment during endometriosis or prevent lesion formation. We also found that lesion CCR2^+^ macrophages were close to CGRP^+^ nociceptors. This indicates that CGRP dictates macrophage recruitment to the PerC during endometriosis and that this CGRP-dependent recruitment surpasses the expected contribution of CCR2/CCL2 or CX_3_CL1/CX_3_CR1 signaling.

Efferocytosis is an important process by which macrophages clear debris to promote resolution and tissue healing (Dalli and Serhan, 2019; Fattori et al., 2020b). We also found that CGRP impaired the efferocytosis of endometriosis cells by macrophages. CGRP also downregulates *Alox15*, an enzyme in the pathway that produces specialized pro-resolving lipid mediators (Chiang and Serhan, 2020; Fattori et al., 2020b). Pro-resolving lipid mediators promote efferocytosis and actively resolve inflammation (Dalli and Serhan, 2019; Fattori et al., 2020b), which could partially explain the CGRP-mediated impairment of efferocytosis. In addition, we found that CGRP promotes endo-epi cell growth by macrophages. CGRP-treated macrophages upregulate the transcripts of chemoattractant molecules such as *Ccl2* (Gregory et al., 2006)*, Ccl13* (Carlsen et al., 2004), and *Il33* (Verri et al., 2010). Furthermore, we found an increase in *Mmp3*, *Mmp8*, *Mmp12,* and *Mmp13*, which are linked to endometrial invasiveness and fibrosis (Ke et al., 2021; Muharam et al., 2023; Stratopoulou et al., 2023). CGRP also increased the chemokines *Cxcl5* (Mao et al., 2020; Walens et al., 2019) and *Col6a1* (Liang et al., 2021), which are responsible for promoting the metastasis and growth of tumors in cancer models. Notably, changes in both efferocytosis capacity and endometrial cell growth stimulation were rescued by rimegepant. Therefore, it is likely that CGRP programs macrophages to a “pro-endometriosis” state that ultimately allows lesion growth and contributes to pain.

In conclusion, we describe TRPV1^+^ nociceptor to macrophage communication via CGRP/RAMP1 signaling that drives the growth of endometriosis lesions and promotes pain. We found that both human and mouse samples contained CGRP^+^ fibers and RAMP1. *In vivo*, we used 4 different FDA approved drugs targeting either CGRP or RAMP1 and found a reduction in pain (evoked and spontaneous) and lesion size. *In vitro*, we found that CGRP programs macrophage to a pro-endometriosis phenotype that is permissive to lesion growth and pain, in a mechanism that is RAMP1-sensitive. Therefore, blocking neuroimmune communication could be a significant and innovative non-hormonal and non-opioid approach for endometriosis treatment.

## Author contributions

Conceptualization: V.F. and M.S.R. Investigation: V.F., T.H.Z., F.S.R.-O., O.K.H., and D.P. Formal analysis: V.F., T.H.Z., F.S.R.-O., O.K.H., A.J, and L.S. Human sample acquisition: M.L.S., A.E.L., and R.M.A. Methodology: V.F., A.J., L.S., and M.S.R. Resources: R.M.A., W.A.V.J., and M.S.R. Project administration: V.F. Supervision: V.F. and M.S.R. Visualization: V.F., T.H.Z., A.J., L.S., W.A.V.J., and M.S.R. Writing (original draft): V.F. Writing (reviewing and editing): all authors. All authors read and approved the final version of the manuscript. Funding acquisition: V.F. and M.S.R.

## Acknowledgments

We thank Neurodevelopmental Behavioral Core (NBC) and Flow Cytometry Research Core of the Boston Children’s Hospital. T.H.Z. thanks the Coordination for the Improvement of Higher Education Personnel (CAPES) for the 12-month scholarship to develop his Split Fellowship (Doutorado Sanduiche PDSE). F.S.R.-O. thanks the National Council for Scientific and Technological Development (CNPq, Brazil) for the 12-month scholarship to develop her Split Fellowship (Doutorado Sanduiche SWE). We also thank Londrina State University’s core facility CMLP-UEL (Central Multiusuário de Laboratórios de Pesquisa da Universidade Estadual de Londrina) for the use of instruments free of charge. T.H.Z. and F.S.R.-O. also acknowledge PhD scholarship from CAPES (finance code 001). W.A.V.J. acknowledges the CNPq Productivity fellowship (307186/2017-2). We also thank Rachel Arredondo for English editing this manuscript.

## Data availability statement

The data that support the findings of this study are available from the corresponding author upon reasonable request.

---

**Fig S1.**
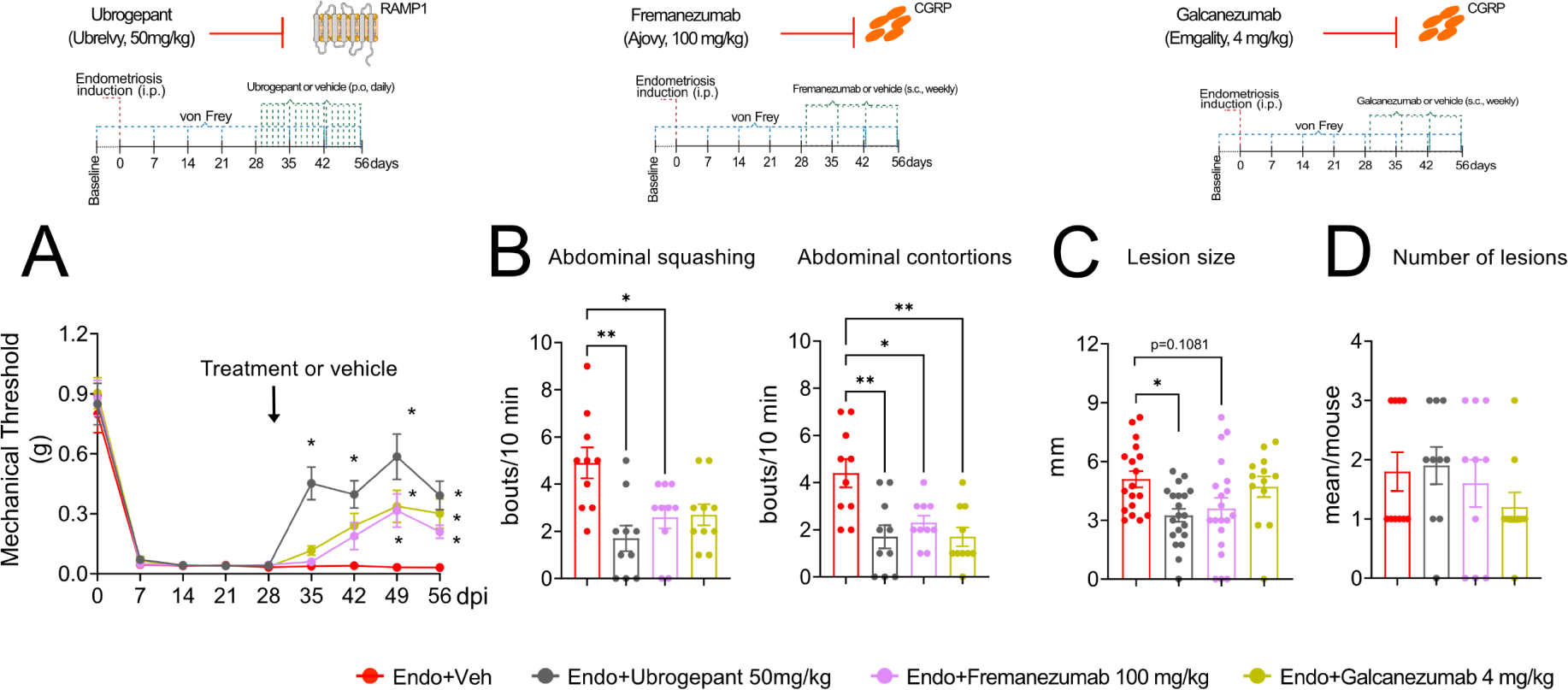
Blocking CGRP/RAMP1 signaling reduces endometriosis-associated pain and lesion size. **(A)** Abdominal mechanical hyperalgesia was measured before (zero) and after (weekly) using von Frey filaments. Results are presented as mean ± SEM of mechanical threshold, n = 10 mice per group (two-way repeated-measures ANOVA followed by Tukey’s post hoc, ****p< 0.0001). **(B)** Spontaneous pain measured by abdominal squashing and abdominal contortions as described in Figure 3 B and G. **(C)** Percentage of mice with lesion was determine by a simple count of visible lesions while **(D)** lesion size was determined in the remaining lesions by measuring perpendicular diameters. Sham mice do not show any lesion (n.d. = not detectable). Results are expressed as mean ± SEM, n = 10 (one-way ANOVA followed by Tukey’s post hoc, ****p< 0.0001; **p<0.02; *p<0.05).

**Fig S2.**
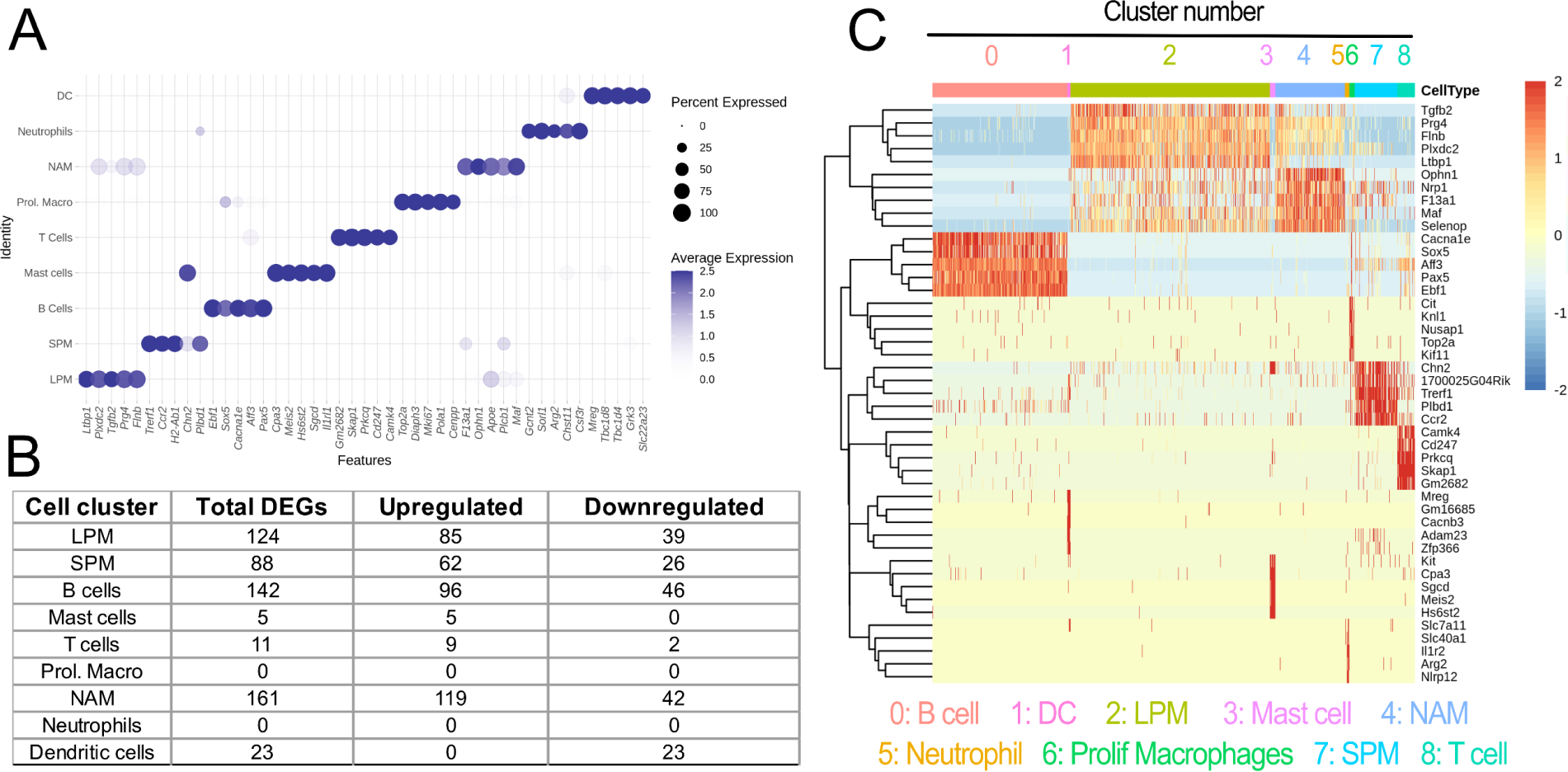
scRNAseq analysis of PerC immune cells during endometriosis. **(A)** Dot plot displaying the top 5 cluster marker genes based on fold-change for the identity of each cell cluster. **(B)** Number of DEGs (total, upregulated, and downregulated) in each cell cluster. **(C)** Heatmap showing normalized expression of 5 top cluster marker genes based on fold-change in immune cells of the PerC cells, with key marker genes highlighted. (n = 5 pooled mouse PerC washes for each condition).

**Fig S3.**
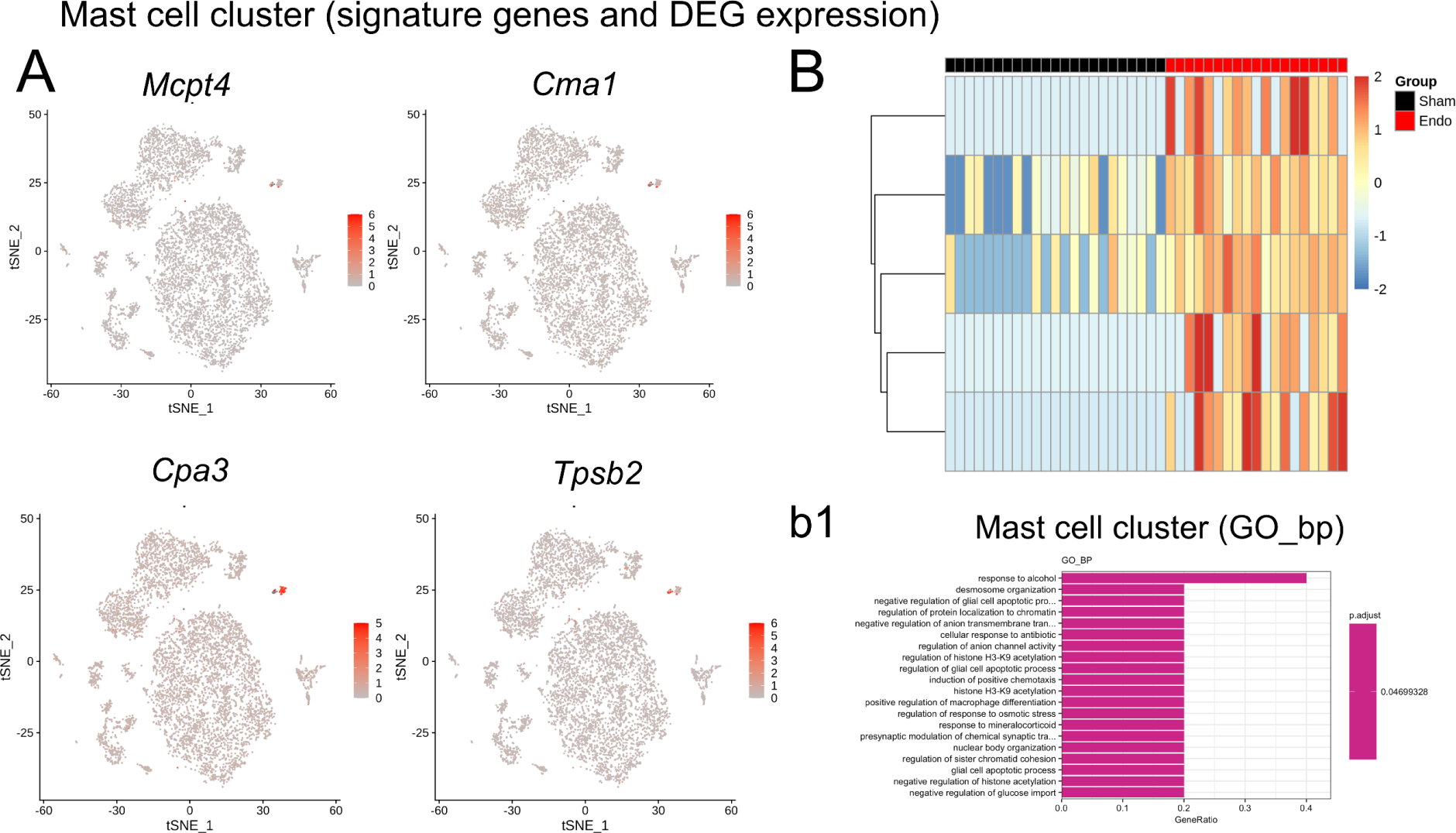
scRNAseq analysis of dendritic cell cluster during endometriosis. **(A)** tSNE plot displaying signature genes for the dendritic cell cluster. **(B)** Heatmap showing DEGs of the dendritic cell cluster in sham vs endometriosis mice. **(b1)** GO enrichment analysis of the DC cluster in sham vs endometriosis mice. (n = 5 pooled mouse PerC washes for each condition).

**Fig S4.**
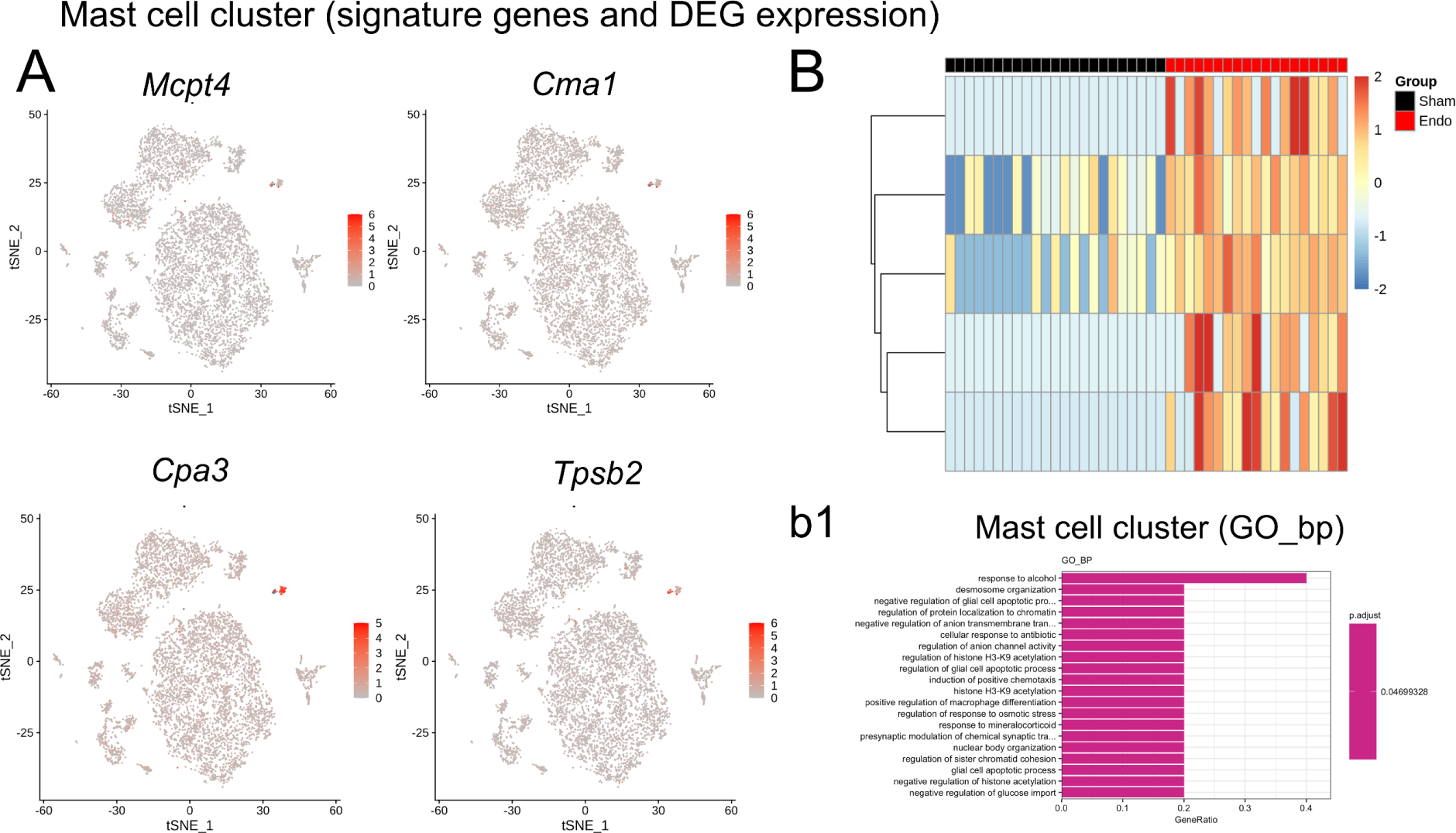
scRNAseq analysis of mast cell cluster during endometriosis. **(A)** tSNE plot displaying signature genes for the mast cell cluster. **(B)** Heatmap showing DEGs of the mast cell cluster in sham vs endometriosis mice. **(b1)** GO enrichment analysis of the mast cell cluster in sham vs endometriosis mice. (n = 5 pooled mouse PerC washes for each condition).

**Fig S5.**
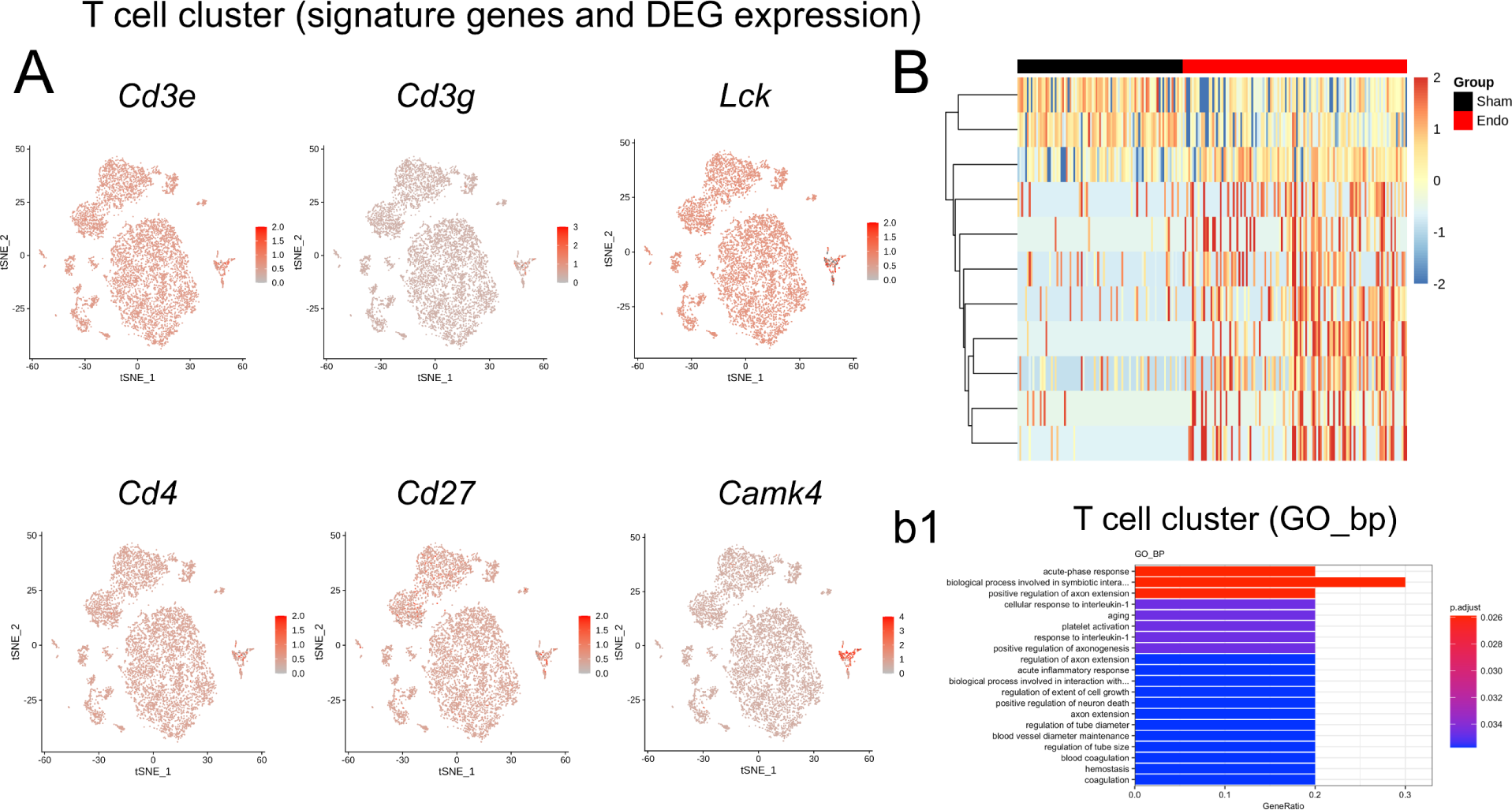
scRNAseq analysis of T cell cluster during endometriosis. **(A)** tSNE plot displaying signature genes for the T cell cluster. **(B)** Heatmap showing DEGs of the T cell cluster in sham vs endometriosis mice. **(b1)** GO enrichment analysis of the T cell cluster in sham vs endometriosis mice. (n = 5 pooled mouse PerC washes for each condition).

**Fig S6.**
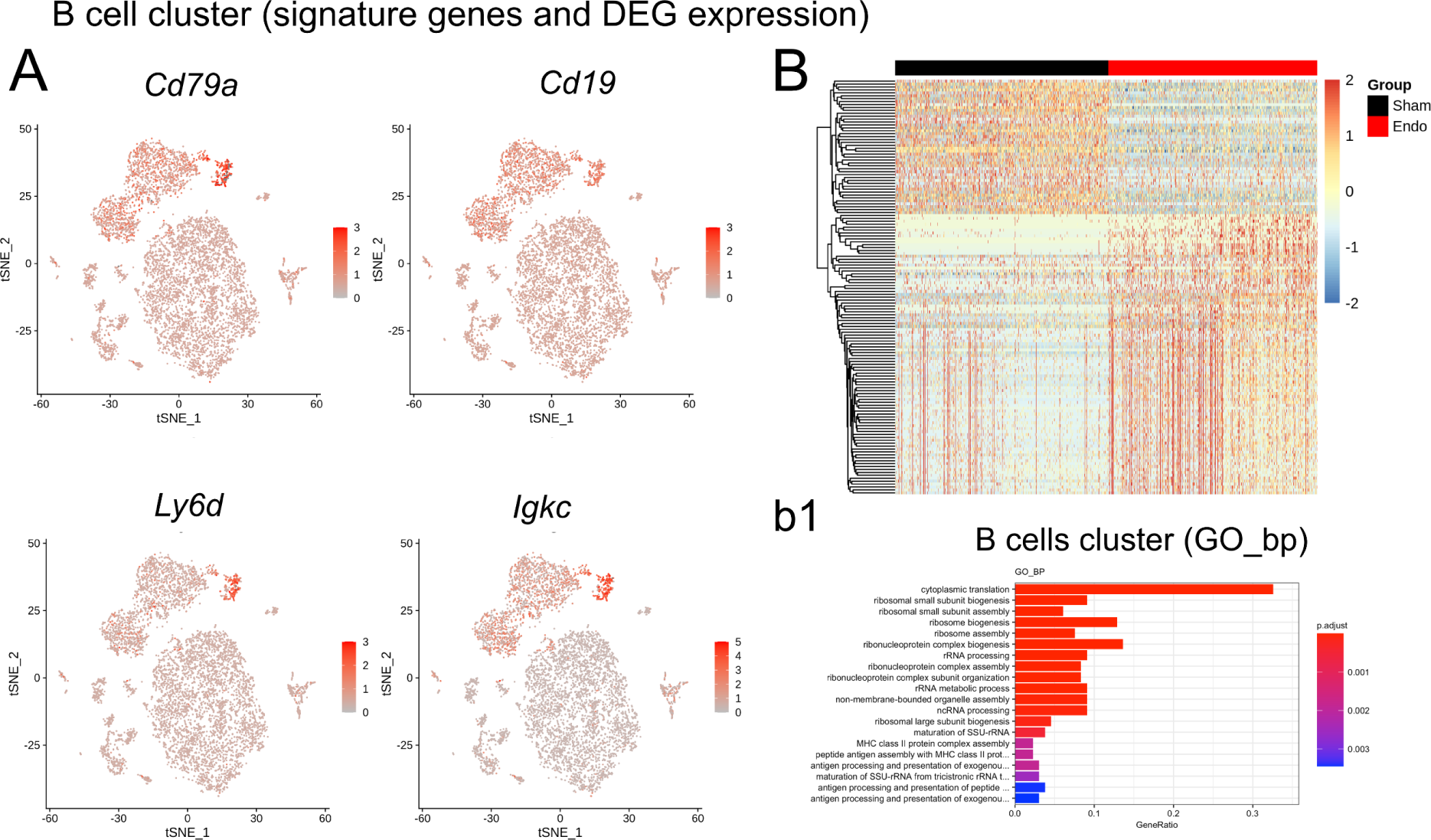
scRNAseq analysis of B cell cluster during endometriosis. **(A)** tSNE plot displaying signature genes for the B cell cluster. **(B)** Heatmap showing DEGs of the B cell cluster in sham vs endometriosis mice. **(b1)** GO enrichment analysis of the B cell cluster in sham vs endometriosis mice. (n = 5 pooled mouse PerC washes for each condition).

**Fig S7.**
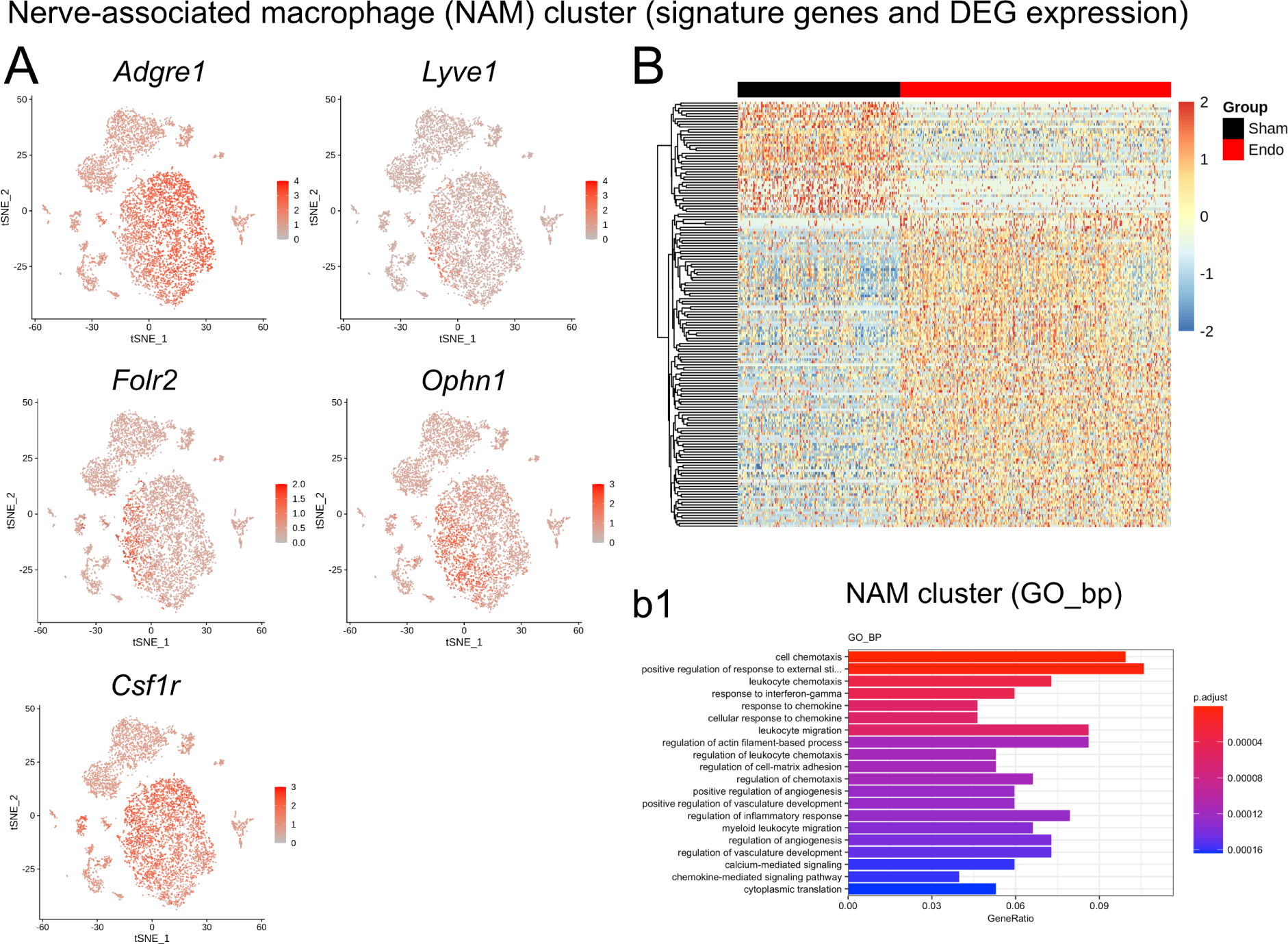
scRNAseq analysis of NAM cluster during endometriosis. **(A)** tSNE plot displaying signature genes for the NAM cell cluster. **(B)** Heatmap showing DEGs of the NAM cell cluster in sham vs endometriosis mice. **(b1)** GO enrichment analysis of the mast cell cluster in sham vs endometriosis mice. (n = 5 pooled mouse PerC washes for each condition).

**Fig S8.**
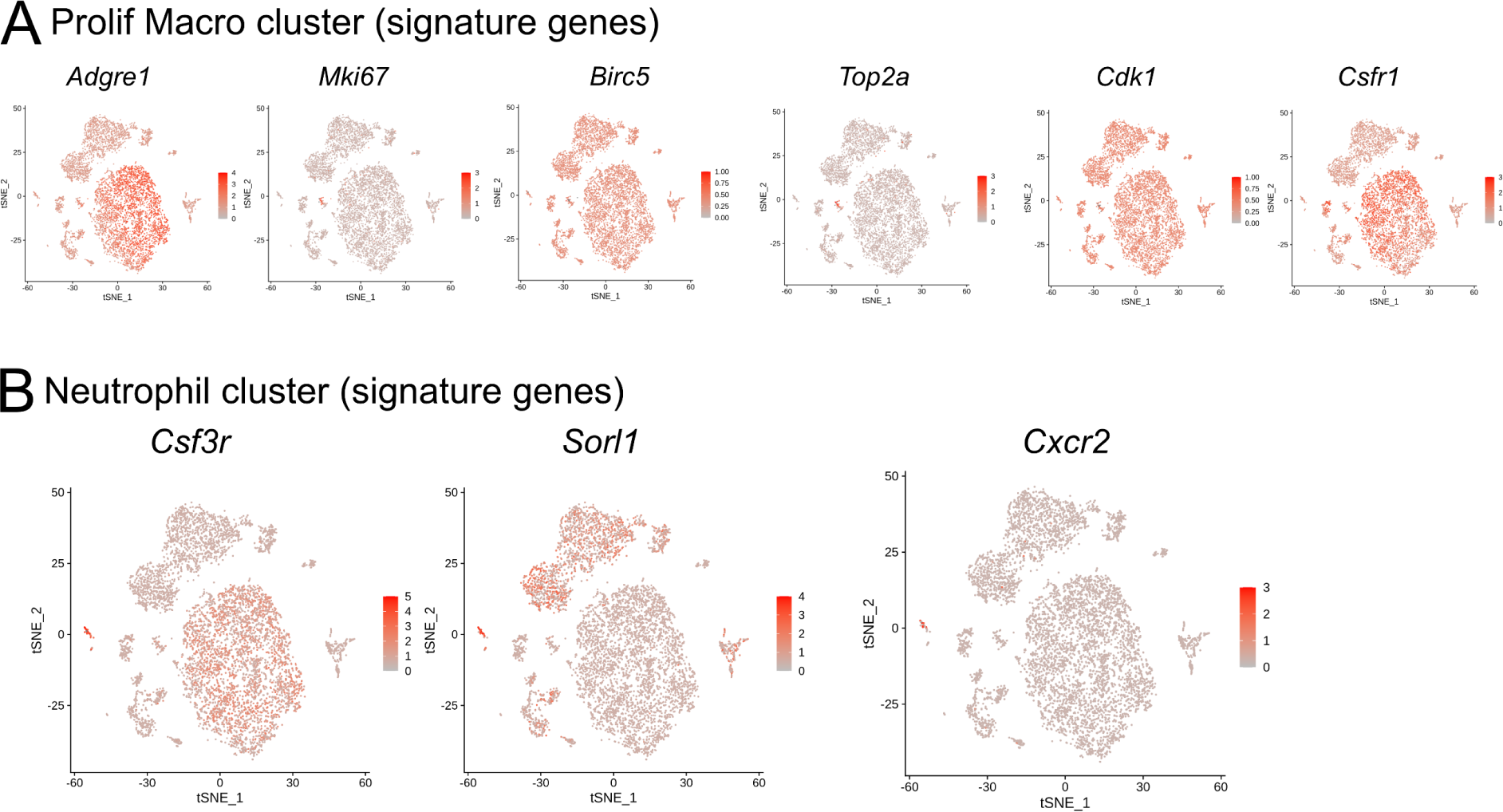
scRNAseq analysis of proliferating (Prolif) macrophage and neutrophil clusters during endometriosis. **(A)** tSNE plot displaying signature genes for the Prolif macrophage cell cluster. **(B)** tSNE plot displaying signature genes for the neutrophil cell cluster. As observed in Fig S2B, no DEGs were found for these two clusters. (n = 5 pooled mouse PerC washes for each condition).

**Fig S9.**
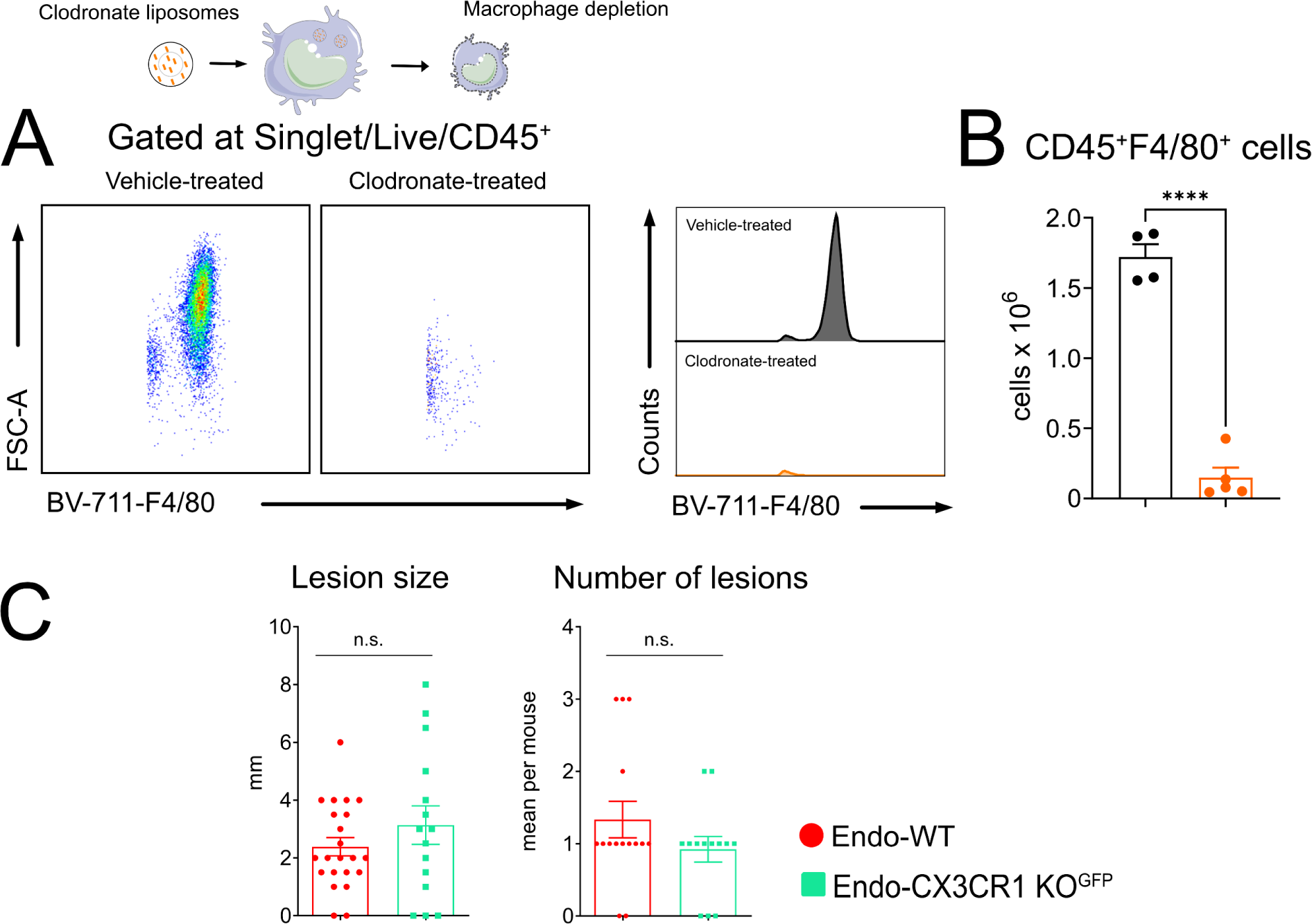
Clodronate depletion of PerC macrophages and CX_3_CR1^GFP^ experiments. **(A)** Flow cytometry data of PerC wash collected from vehicle- or clodronate (0.0625 µg per mouse)-treated mice. Macrophages were gated as CD45^+^F4/80^+^ cells. Panel **B** shows cell quantification. Results are presented as mean ± SEM. n = 4 (vehicle liposomes) or 5 (clodronate liposomes) mice per group. To determine the role of CX_3_CL1 in endometriosis, we used CX_3_CR1^GFP^ mice. **(C, left)** Lesion size was determined in the remaining lesions by measuring height and weight. **(C, right)** Number of visible lesions was measured as sum of total lesions per mouse. Results are expressed as mean ± SEM, n = 15 (WT) and 13 (CX_3_CR1^GFP^) (Student’s t-test. ****p< 0.0001).

