## Supplementary material for "Nociceptor to macrophage communication through CGRP/RAMP1 signaling drives endometriosis-associated pain and lesion growth": Material and methods

### 1. Material and methods

#### *1.1 Animals*

All procedures were performed according to the National Institutes of Health Guide for the Care and Use of Laboratory Animals, International Association for the Study of Pain (IASP), as well as were in accordance with the laws of the United States and regulations of the Department of Agriculture. All experiments were approved by the Institutional Animal Care and Use Committee (IACUC) at Boston Children's Hospital (protocols #19-12-4054R and #00001816) and Londrina State University (protocols #2669.2019.54 and #047.2023). Healthy female C57BL/6 mice (Stock #000664), B6.129-Trpv1$^{tm1(cre)Bbm}$/J (TRPV1cre, sotck #017769), and B6.129P2-Gt(ROSA)26$^{Sortm1(DTA)Lky}$/J, a Cre-dependent diphtheria toxin A (DTA) strain (ROSA-DTA, stock #009669) were purchased from Jackson Laboratories (Bar Harbor, ME, USA) and experiments were performed at Boston Children's Hospital (Boston, MA, USA). B6.129S4-Ccr2$^{tm1Ifc}$/J (CCR2-KO, stock #004999), B6.129(Cg)-Ccr2$^{tm2.1Ifc}$/J (CCR2-KO$^{RFP}$, stock #017586), and B6.129P2(Cg)-Cx3cr1$^{tm1Litt/}$J (CX$_3$CR1-KO$^{GFP}$, stock #005582) were received from Ribeirão Preto Medical School (University of São Paulo, Ribeirão Preto, SP, Brazil) and experiments performed at Londrina State University (Londrina, PR, Brazil). Female age-matched mice from 8 to 14 weeks of age were used for all experiments in this study. Mice were randomly assigned and housed in standard clear plastic cages with no more than 5 mice per cage in a 12:12h light/dark cycle with *ad libitum* access to water and food. Behavioral testing was performed between 9 a.m. and 5 p.m. in a room maintained at a temperature of 21±1°C. All efforts were made to minimize the number of animals used and their suffering. Euthanasia was performed by controlled $CO_2$ inhalation.

#### *1.2 Human sample collection and demographics*

Samples from six patients undergoing laparoscopic excision/ablation of endometriosis lesions (ectopic lesion or endometrioma cases) were collected and divided by the Dr. Raymond Anchan into two parts in accordance with Institutional Review Board (IRB) approval (Mass General Brigham IRB #2017P002765). Half of each sample was sent to histopathological confirmation of endometriosis and the other half was taken to the lab for processing. All patients were recruited at the Brigham Women's Hospital (Boston, MA, USA), and presented clinical and histopathological features that fulfilled the criteria for endometriosis. All patients were informed about the aims of the study and provided written consent before participating. Pain score was determined using the Endometriosis Phenome and Biobanking Harmonization Project (EPHect) protocol (Rahmioglu et al., 2014). Patient demographics and pain score are presented in Table 1.

Table 1. Endometriosis patient demographics.

| **Age** | **Medication** | **Cycling (Y or N)** | **Age of symptom onset** | **Disease stage** | **Lesion color** | **Pain score** |
|---|---|---|---|---|---|---|
| 29 | Alyacen | Y | 25 | 4 | Mostly black, white, and red | 6 |
| 41 | Aygestin | Y | 13 | 4 | Black | 8 |
| 26 | None | Y | 19 | 3 | Red | 10 |
| 42 | Aygestin | Y | 30 | 3 | Black | 4 |
| 44 | Ferrous Sulfate, Cephalexin | Y | 38 | 1 | Red | Mild pain* |
| 35 | Ferrous Sulfate | Y | 33 | 2 | Filmy | 10 |

*Pain was self-reported by the patient to Dr. Anchan.

*1.3 Induction of endometriosis*

After at least one week of acclimatization, donor mice received a subcutaneous injection of 3 µg/mouse estradiol benzoate (30 µg/mL in sesame oil) to stimulate the growth of the endometrium as previously described (Fattori et al., 2020). Four days later, the uteri of the donor mice were dissected into a Petri dish containing sterile Hank's Balanced Salt Solution (HBSS, Thermo Fisher Scientific, Waltham, MA, USA), and minced with scissors and scalpel one at the time, ensuring that the maximal diameter of each fragment was consistently smaller than 1 millimeter (mm). Each dissociated uterine horn was then injected intraperitoneally into a recipient mouse in 500 µL of sterile HBSS, meaning that one uterus of a donor mouse was used for endometriosis induction in every two recipient mice. Sham mice received an intraperitoneal injection of 500 µL of sterile HBSS.

*1.4 Mouse sample immunofluorescence*

*1.4.1 Sample preparation*

Samples (lesions, PerC wash, or DRG neurons) were dissected at 28 dpi for immunofluorescence. Lesions or paired thoracic and lumbosacral (T10-L3 plus L6-S1) dorsal root ganglia (DRG) were dissected at and maintained in 4% paraformaldehyde (PFA, for 24h) and then to 30% sucrose (for 72h). PerC wash was collected into FACS buffer (PBS, 0.5% BSA, and 2.5 mM EDTA) and cells were centrifuged for 10 min at 300g. The resulting supernatant was discarded, and the pellet resuspended in 500 µL of FACS buffer. For lesions and DRG neurons, optimum cutting temperature reagent (Tissue-Plus O.C.T., Fisher Healthcare, Thermo Fisher Scientific, Waltham, MA, USA)-embedded samples were cut in 16 µm sections while PerC wash cells were mounted on slides with Fluoromount-G compound (cat #0100-01, SouthernBiotech, Birmingham, AL, USA).

*1.4.2 Staining*

Before incubation with antibodies, slides were blocked with 5% goat serum in PBS, 0.3% Triton X-100 (0.3% PBST). Primary antibodies used in this study are as follow: anti-PGP9.5 (1:50, cat #ab8189, Abcam, Cambridge, MA, USA), anti-CGRP (1:500, cat #C8198, MilliporeSigma, Burlington, MA, USA); anti-RAMP1 (1:50, cat # PA5-110265, Life Technologies, Thermo Fisher Scientific, Waltham, MA, USA); anti-TRPV1 (1:250, cat #ACC-030; Alomone Labs, Jerusalem, Israel), anti-F4/80 (1:500, cat #14-4801-82, Life Technologies, Thermo Fisher Scientific, Waltham, MA, USA) anti-pNF-κB p65 (1:50, cat #sc-166748, Santa Cruz Biotechnology, Dallas, TX, USA); anti-Ly6C (1:50, cat #sc-271811, Santa Cruz Biotechnology, Dallas, TX, USA); phospho-CREB (1:350, clone 87G3, cat #9198S, Cell Signaling Technology, Beverly, MA, USA). All primary antibodies and DAPI (2 µg/ml; 1:500, cat #62248, Life Technologies, Thermo Fisher Scientific, Waltham, MA, USA) were incubated overnight at 4 °C (2.5% goat serum in 0.3% PBST) and then washed four times with 0.3% PBST before proceeding with secondary antibody incubations. Secondary antibodies used were: goat anti-mouse Alexa 488 (1:4000, cat #A-10680, Life Technologies, Thermo Fisher Scientific, Waltham, MA, USA); goat anti-rabbit Alexa Fluor 488 (1:1000, cat #A-11029, Life Technologies, Thermo Fisher Scientific, Waltham, MA, USA); goat anti-rat Alexa Fluor 647 (1:500, cat #A-21247, Life Technologies, Thermo Fisher Scientific, Waltham, MA, USA); goat anti-mouse Alexa Fluor 647 (1:500, cat #A-21236, Life Technologies, Thermo Fisher Scientific, Waltham, MA, USA); goat anti-rabbit DyLight 488 (1:200, cat #DI-

1488, Peterborough, United Kingdom); goat anti-rabbit DyLight 594 (1:200, cat #DI-1594, Peterborough, United Kingdom) and incubated for 50 min at room temperature (2.5% goat serum in 0.3% PBST). Stained slides were then washed five times with 0.3% PBST and mounted in Fluoromount-G compound. Images were taken and processed on Zeiss LSM 880 laser scanning microscope using 20x objective (Carl Zeiss Microscopy, Thornwood, NY, USA). For experiments involving CCR2-KO$^{RFP}$ mice, the images were taken on a TCS SP8 confocal microscope (Leica Microsystems, Mannheim, Germany). Fluorescence intensity was measured using ImageJ (NIH, Bethesda, MD, USA).

*1.5 Human sample immunofluorescence*

*1.51. Sample preparation*

Samples were rinsed in PBS without calcium or magnesium, cut into smaller sections, and placed into 4% PFA for 2h at room temperature. Samples were then added to increasing percentages of sucrose (10% and 20% sucrose at room temperature) for 2h each, and then left in 30% overnight at 4 °C. The next day samples were embedded in OCT (Fisher Healthcare, Thermo Fisher Scientific, Waltham, MA, USA) and kept in -80 °C for long-term storage until sectioning and slide preparation. OCT-embedded samples were cut in 16 µm sections.

*1.5.2 Staining*

Before incubation with antibodies, slides were blocked with 5% goat serum in PBS, 0.3% Triton X-100 (0.3% PBST). Primary antibodies used were anti-CGRP (1:100, cat #RA24112; Neuromics, Edina, MN, USA) and anti-RAMP1 (1:50, cat #PA5-110265, Life Technologies, Thermo Fisher Scientific, Waltham, MA, USA). All primary antibodies and DAPI (2 µg/ml; 1:500, cat #62248, Life Technologies, Thermo Fisher Scientific, Waltham, MA, USA) were incubated overnight at 4 °C (2.5% goat serum in 0.3% PBST) and then washed four times with 0.3% PBST before proceeding with secondary antibody incubations. Secondary antibody was goat anti-rabbit DyLight 594 (1:200, cat #DI-1594, Peterborough, United Kingdom) and incubated for 50 min at room temperature (2.5% goat serum in 0.3% PBST). Stained slides were then washed five times with 0.3% PBST and then covered with Sudan Black B (0.05% diluted in 70% Ethanol), a blocker of lipofuscin, for 10 min. Slides were washed three additional times with 0.3% PBST and then mounted in Fluoromount-G compound (cat #0100-01, SouthernBiotech, Birmingham, AL, USA). Images were taken and processed on a Zeiss LSM 880 laser scanning microscope using 20x objective (Carl Zeiss Microscopy, Thornwood, NY, USA). Fluorescence intensity measured using ImageJ (NIH, Bethesda, MD, USA).

*1.6 Flow cytometry*

Peritoneal cavity wash was collected into fluorescence-activated cell sorting (FACS) buffer (PBS, 0.5% BSA, and 2.5 mM EDTA) and cells were centrifuged for 10 min at 300g. The resulting supernatant was discarded, and the pellet resuspended in 100 µL of FACS buffer. The cell suspension was incubated on ice with mouse FcR Blocking Reagent (cat #130-092-575, Miltenyi Biotec, Cambridge, MA, USA) for 10 min, washed and then centrifuged for 10 min at 300g. The resultant pellet was then resuspended and incubated for 30 min on ice with the following antibodies (BioLegend, San Diego, CA, USA): anti-CD45-AlexaFluor® 488 (1:200, clone 30-F11, cat #103122), anti-F4/80-Brilliant Violet 711™ (1:50, clone BM8, cat #123147), and anti-Ly6C-PE/Dazzle™ 594 (1:200, clone HK1.4, cat #128044). Live cell dye was Invitrogen™ eBioscience™ Fixable Viability Dye eFluor™ 450 (1:1000, cat #50-169-62, Thermo Fisher Scientific, Waltham, MA, USA). Cells were then washed and centrifuged for 10 min at 300g, and the pellet resuspended in 300 µL of 2% PFA in FACS buffer. Flow cytometry was performed on a LSR II flow cytometer (BD Biosciences) and data analyzed and plotted using FlowJo software (FlowJo LLC).

*1.7 Calcium imaging*

Calcium imaging of DRG neurons was performed as previously described (Fattori et al., 2020). These ganglia were chosen because of their involvement in abdominal pain and pelvic organ cross-sensitization (Christianson et al., 2007; Christianson et al., 2006; Malykhina et al., 2006). DRGs were dissected into Neurobasal-A medium (Life Technologies, Thermo Fisher Scientific, Waltham, MA, USA), dissociated in collagenase A (1 mg/mL)/dispase II (2.4 U/mL) (RocheApplied Sciences, Indianapolis, IN, USA) in HEPES-buffered saline (MilliporeSigma, Burlington, MA, USA) for 70 min at 37°C. After trituration with glass Pasteur pipettes of decreasing size, DRG cells were centrifuged over a 10% BSA gradient, plated on laminin-coated cell culture dishes. DRGs were loaded with 5 uM of Fura-2AM in Neurobasal-A medium, incubated for 30 min 37°C, washed with HBSS and imaged in using an Eclipse Ti-S/L100 inverted microscope (Nikon Instruments, Melville, NY, USA). An ultraviolet light source (Lambda XL lamp, Sutter Instrument) was used for excitation of Fura-2-AM by alternating 340 nm and 380 nm wavelengths. NIS-elements software (Nikon Instruments, Melville, NY, USA) was used to image, process, and analyze 340/380 ratiometric images from DRG neurons. An increase in 340/380 ratio of 10% or more from baseline levels was considered a positive response to a ligand. DRG plates were recorded for 12 min, which was divided in: two minutes of initial reading (0 second mark, baseline values), following by stimulation with the filtered (0.22µM) homogenate of endometriosis lesion for four minutes at 120 second mark, and then capsaicin (500 nM, TRPV1 agonist) or AITC (100 µM, TRPA1 agonist) for four min at 360 seconds mark and KCl for two min at 600 seconds mark (40 mM, activates all neurons).

*1.8 Behavioral testing*

For mechanical and heat hyperalgesia tests, mice were allowed to habituate to the apparatus for at least 2h and during three consecutive days before the beginning of measurements as previously described (Fattori et al., 2020). After habituation, baseline measurements were obtained on two consecutive days prior to induction of endometriosis. Pain intensity to a mechanical stimulus (mechanical hyperalgesia) in the abdominal region was measured using von Frey filaments. The experimenter was trained, and care was taken not to stimulate the same point consecutively and the stimulation of the external genitalia was avoided. One of the following behaviors was considered as a withdrawal response: jump or paw flinches (Fattori et al., 2020). The mechanical threshold was determined by the up and down method starting with 0.4g filament and calculated using the open source software Up-Down Reader (Gonzalez-Cano et al., 2018).

For spontaneous abdominal pain measurements, stretching the abdomen (abdominal contortions) and squashing of the lower abdomen against the floor were quantified as previously described (Fattori et al., 2020). Briefly, for abdominal contortions, mice were placed in individual chambers in a temperature-controlled (29 °C) glass plate and the number of abdominal contortions was quantified for 10 min. Positive responses consisted of a contraction of the abdominal muscle together with stretching of the hind limbs. For abdominal squashing, the number of times the mice pressed the lower abdominal region against the floor in 10 minutes was quantified. When treatment was included, vehicle- or drug-treated mice received the treatment starting at 29 dpi. In all testing, the investigators were blinded to the treatments.

*1.9 Cytokine and chemokine level measurements*

Lesions were dissected into lysis buffer (PBS, 1% Triton X-100 and protease inhibitors) and then homogenized and centrifuged (10,000 g × 5 min, 4 ◦C). The resulting supernatants were used to determine protein levels using Pierce™ BCA Protein Assay Kit (Life Technologies, Thermo Fisher Scientific, Waltham, MA, USA) and the cytokine array. The assay was conducted with 800 µg of protein for each group for the Proteome Profiler Mouse XL Cytokine Array (cat #ARY028) and 600 µg of protein for each group for the Proteome Profiler Mouse Angiogenesis Array (cat #ARY015) kits. One membrane was used for each group and each membrane corresponded to the protein levels of a pool of 5 lesions or 5 donor uterus per group. The analysis of each membrane was conducted by densitometry using Gilles Carpentier's Dot-Blot-Analyzer macro with the values corrected by the mean values of the reference spots as previously described (Fattori et al., 2022). The macro was written for ImageJ software (NIH, Bethesda, MD, USA) and is available at https://imagej.nih.gov/ij/macros/toolsets/Protein%20Array%20Analyzer.txt and more information can be found at http://image.bio.methods.free.fr/dotblot.html. Since sham mice do not develop lesions, uterine horns from donor mice were used to determine baseline levels of each cytokine. CCL2 (cat #MJE00B), CCL3 (cat #MM00), $CX_3CL1$ (cat #MCX310), PLGF (cat #MP200), and VEGF (cat #MMV00) levels were determined by enzyme-linked immunosorbent assay (ELISA) kits. All ELISA and array kits were purchased from R&D Systems (Minneapolis, MN, USA).

*1.10 Lesion size and number of lesions*

At the specified timepoints, lesions were carefully dissected and measured using a caliper. Lesion size is expressed in millimeters (mm) calculated from the mean of two measurements (width and height) while number of lesions per mouse was determined by a simple count of the visible lesions (Fattori et al., 2020)

*1.11 Single cell RNA sequencing*

Peritoneal cavity washes from sham littermate controls and lesion-bearing TRPV1-creDTA and littermate control mice were collected at 28dpi and used for droplet-based scRNA-seq (10x Genomics, Pleasanton, CA, USA). Library preparation was conducted using the Next GEM Single Cell 3′ Reagent Kits v3.1 Dual Index Kit TT, Set A (10x Genomics) following the manufacturer's instructions. Sequencing using an Illumina NovaSeq 6000 System was conducted at Novogene, Inc. The quality of the single-cell suspensions (viability >96%, concentration = 1,200 cells/μL) was confirmed immediately before encapsulation. Encapsulation of cells was performed in a Chromium Controller (10x Genomics, Pleasanton, CA, USA) targeting 4,000 cells per sample. Following encapsulation and mRNA barcoding, cDNA was synthesized, isolated and amplified (12 cycles) using a Single Cell 3′ GEM kit (10x Genomics) and a SPRIselect reagent kit (Beckman Coulter). The quality of the amplified cDNA (concentration, size, and purity) was verified using a High Sensitivity D5000 ScreenTape and 4200 TapeStation system (Agilent Technologies, Santa Clara, CA, USA). Next, amplified cDNA was used for library construction. cDNA fragmentation, end repair, A-tailing, adaptor ligation and sample index PCR amplification were performed using Chromium Next GEM Single Cell 3′ Reagent Kits v3.1 Dual Index Kit TT, Set A (cat# PN3000431, 10x Genomics). After library construction, quality control was performed using a High Sensitivity D5000 ScreenTape and 4200 TapeStation system (Agilent Technologies). For sequencing, sample libraries were pooled and equally distributed across the lanes of an Illumina NovaSeq S1 flow cell (Read 1: 28 bp; i5 and i7 indexes: 10 bp; Read 2: 90 bp).

The raw single-cell RNA sequencing data was preprocessed using CellRanger v7.1.0 (10x Genomics), including aligning reads to the mouse mm10 reference genome, and generating cellxgene expression count matrices. The generated expression count matrices were processed and analyzed using an in-house single-cell RNA-seq pipeline. The pipeline was built based on the Seurat R package (R v4.2.3, Seurat v4.3.0), including ambient RNA removal, quality control, cell filtering, spectral clustering, cell type annotation, differential gene expression, and visualization (Butler et al., 2018; Stuart et al., 2019). Ambient RNA was removed using SoupX (v.1.6.2) (Young and Behjati, 2020). Multiplets were identified using the scds package (v.1.14.0) using the cxds function (Bais et al, 2020). Only identified singlets were kept for further analysis. Cells with extreme (very low or very high, out of the 95% CI) library sizes, number of features and high content of mitochondrial reads (>10%) were filtered out. Cells of good quality from the two replicates per group were merged, and the three experimental groups were integrated for analysis using Harmony (Korsunsky et al., 2019). Principal component analysis (PCA) over the identified 2000 highly variable genes was applied for data dimension reduction (dimensions = 30) prior to cell clustering. Cell clustering was performed on integrated data with a shared nearest-neighbor (KNN) graph-based method using the FindNeighbors function included in Seurat, followed by the Louvain algorithm for modularity optimization (resolution = 0.5) using FindClusters function. After the cell clusters were determined, their top marker genes were identified with the FindMarkers function. For cluster annotation, the top marker genes based on the adjusted p-value were manually curated to match canonical cell types and their marker genes based on literature research. Differentially expressed genes (DEGs) between the conditions for each cell type were identified by using the Seurat FindMarkers function (adjusted p-value < 0.05). Gene ontology (GO) enrichment tests was performed by the clusterProfiler R package (Yu et al., 2012). A GO term was treated as significantly enriched if an adjusted p-value (with Benjamini-Hochberg correction) is <0.05. All bar plots illustrating significant pathway or GO terms were created using the enrichplot R package (Yu et al., 2019).

*1.12 Cell culture*

Peritoneal cavity wash was collected in FACS buffer from the abdominal cavity of naïve C57BL/6 mice. Peritoneal cavity immune cells were seeded in petri dishes in DMEM-F12 media and incubated at 37 °C in 5% $CO_2$ for 60 minutes. Non-adherent cells were discarded, and adherent cells (macrophages) were scraped, centrifuged, and counted. For experiments involving mouse endometrial epithelial cells (endo-epi cells) cell proliferation, macrophages were plated (100,000 cells per well) onto tissue culture insert and left overnight with 20,000 endo-epi cells plated on the bottom of the well. Stimulation with CGRP (100 nM, cat #RP11095, Genscript, Piscataway, NJ, USA) or vehicle was performed for 24 hours, and then endo-epi cells were used to determine cell proliferation assay using a CyQuant™ kit (cat #C7026, Life Technologies, Thermo Fisher Scientific, Waltham, MA, USA) in a BioTek Synergy H1 multimode reader (Agilent, Santa Clara, CA, USA). Treatment with rimegepant (100 nM, cat #HY-15498, MedChemExpress LLC, Monmouth Junction, NJ, USA) or vehicle was performed 30 minutes before stimulus with CGRP. For the efferocytosis assay, macrophages (200,000 cells per well) were plated overnight onto a 96-

well plate. Stimulation with CGRP (100 nM) or vehicle was performed for 24 hours. After that, etoposide-induced apoptotic 12Z-eGFP were added (600,000 cells per well) to the macrophage culture and incubated for 1 hour. Cells were washed 3 times with DPBS to remove non-engulfed 12Z-eGFP and then and fluorescence intensity determined using a BioTek Synergy H1 multimode reader. Treatment with rimegepant (100 nM) or vehicle was performed 30 minutes before stimulus with CGRP.

*1.13 Bulk RNA sequencing*

Peritoneal cavity wash was collected in FACS buffer from the abdominal cavity of naïve C57BL/6 mice. Peritoneal cavity immune cells were seeded in petri dishes in DMEM-F12 media and incubated at 37 °C in 5% $CO_2$ for 60 minutes. Non-adherent cells were discarded, and adherent cells (macrophages) were scraped, centrifuged, and counted. Macrophages ($1x10^6$ cells) and mouse endometrial epithelial cell (endo-epi cells, $2x10^5$ cells) were then co-cultured. Stimulation with CGRP (100 nM) or vehicle was performed for 24 hours. After that, conditioned media was collected, filtered, and stored at -80 °C until use. For bulk RNAseq analysis, macrophages ($1x10^6$ cells/flask) were stimulated with conditioned media from vehicle- or CGRP-treated co-cultured experiments for 24 hours. Cells (viability >98%) were then harvested and RNA was extracted using PureLink™ RNA Mini Kit (cat #12183025, Life Technologies, Thermo Fisher Scientific, Waltham, MA, USA). Sequencing was conducted at Novogene, Inc. Differential gene expression analysis was performed in sequenced datasets using the R package DESeq2. Genes were considered differentially expressed when the adjusted p-value was <0.05. The list of differentially expressed genes was used in pathway enrichment analysis and to create volcano plots.

*1.14 Statistical analysis*

Results are presented as mean ± SEM. Data were analyzed using the software GraphPad Prism version 9.2 (GraphPad Software, San Diego, CA, USA). Two-way repeated measure analysis of variance (ANOVA), followed by Tukey's *post hoc*, was used to analyze data from experiments of multiple time points (von Frey). One-way ANOVA followed by Tukey's *post hoc* was used to analyze data from experiments of single time point. Comparison between two groups was conducted using Student's t-test. For the percentage of mice with visible lesions, statistical analysis was estimated by the Kaplan-Meier method followed by the log rank. For all analysis, statistical differences were considered significant when $p < 0.05$.
